# Targeting *de novo* lipogenesis and the Lands cycle induces ferroptosis in KRAS-mutant lung cancer

**DOI:** 10.1101/2021.03.18.434804

**Authors:** Caterina Bartolacci, Cristina Andreani, Gonçalo Vias Do Vale, Stefano Berto, Margherita Melegari, Anna C. Crouch, Dodge L. Baluya, George Kemble, Kurt Hodges, Jacqueline Starrett, Katerina Politi, Sandra L. Starnes, Daniele Lorenzini, Maria Gabriela Raso, Luisa Solis Soto, Carmen Behrens, Humam Kadara, Boning Gao, David Gerber, Ignacio I. Wistuba, John D. Minna, Jeffrey McDonald, Pier Paolo Scaglioni

**Affiliations:** Department of Internal Medicine, University of Cincinnati College of Medicine, Cincinnati, OH 45219, USA; McDermott Center for Human Growth and Development, The University of Texas Southwestern Medical Center, Dallas, TX 75390, USA; Department of Neuroscience, The University of Texas Southwestern Medical Center, Dallas, TX 75390, USA; Department of Interventional Radiology, The University of Texas MD Anderson Cancer Center; Tissue Imaging and Proteomics Laboratory, Washington State University, Pullman, WA 99164.; Sagimet Biosciences, San Mateo, CA 94402; Department of Pathology, University of Cincinnati College of Medicine, Cincinnati, OH 45219, USA; Yale Cancer Center, Yale School of Medicine, New Haven, Connecticut; Department of Surgery, Division of Thoracic Surgery, University of Cincinnati College of Medicine, Cincinnati, OH 45219, USA; Department of Pathology, Fondazione IRCCS Istituto Nazionale dei Tumori di Milano, via Venezian 1, 20133 Milan, Italy; Department of Translational Molecular Pathology; Department of Thoracic H&N Med Oncology, The University of Texas MD Anderson Cancer Center.; Hamon Center for Therapeutic Oncology Research, The University of Texas Southwestern Medical Center, Dallas, TX 75390, USA

**Author notes:** These authors contributed equally to this manuscript (Caterina Bartolacci, Cristina Andreani). Corresponding author: Pier Paolo Scaglioni, The Vontz Center for Molecular Studies, 3125 Eden Avenue, Rm. 3118, Cincinnati, OH 45219-2293.

## Abstract

Mutant *KRAS* (KM) is the most common oncogene in lung cancer (LC). KM regulates several metabolic networks, but their role in tumorigenesis is still not sufficiently characterized to be exploited in cancer therapy. To identify metabolic networks specifically deregulated in KMLC, we characterized the lipidome of genetically engineered LC mice, cell lines, patient derived xenografts and primary human samples. We also determined that KMLC, but not EGFR-mutant (EGFR-MUT) LC, is enriched in triacylglycerides (TAG) and phosphatidylcholines (PC). We also found that KM upregulates fatty acid synthase (FASN), a rate-limiting enzyme in fatty acid (FA) synthesis promoting the synthesis of palmitate and PC. We determined that FASN is specifically required for the viability of KMLC, but not of LC harboring EGFR-MUT or wild type KRAS. Functional experiments revealed that FASN inhibition leads to ferroptosis, a reactive oxygen species (ROS)-and iron-dependent cell death. Consistently, lipidomic analysis demonstrated that FASN inhibition in KMLC leads to accumulation of PC with polyunsaturated FA (PUFA) chains, which are the substrate of ferroptosis. Integrating lipidomic, transcriptome and functional analyses, we demonstrated that FASN provides saturated (SFA) and monounsaturated FA (MUFA) that feed the Lands cycle, the main process remodeling oxidized phospholipids (PL), such as PC. Accordingly, either inhibition of FASN or suppression of the Lands cycle enzymes PLA2 and LPCAT3, promotes the intracellular accumulation of lipid peroxides and ferroptosis in KMLC both *in vitro* and *in vivo*. Our work supports a model whereby the high oxidative stress caused by KM dictates a dependency on newly synthesized FA to repair oxidated phospholipids, establishing a targetable vulnerability. These results connect KM oncogenic signaling, FASN induction and ferroptosis, indicating that FASN inhibitors already in clinical trial in KMLC patients (NCT03808558) may be rapidly deployed as therapy for KMLC.

## Introduction

Mutant *KRAS* (KM) lung cancer (LC) is associated with poor prognosis and resistance to therapy. *KM* expression is not only sufficient to initiate LC but also essential for the viability of KMLC (Fisher et al., 2001; Sunaga et al., 2011). Notwithstanding the notable exception of the *KRAS^G12C^* mutant (Canon et al., 2019; Hallin et al., 2020; Janes et al., 2018; McCormick, 2019), KM remains undruggable. Furthermore, targeting the KM signaling network has proved to be either ineffective or toxic (Tomasini et al., 2016). Finally, cancer immunotherapy benefits only a minority of KMLC patients (Carbone et al., 2017; Reck et al., 2016). Therefore, there is an urgent need for novel therapeutic strategies for KMLC.

KM reprograms cellular metabolism, promoting aerobic glycolysis and lipid synthesis/uptake (Boroughs and DeBerardinis, 2015; Kamphorst et al., 2013a; Padanad et al., 2016). Notably, KM regulates the expression of fatty acid synthase (FASN), the rate-limiting enzyme of *de novo* lipogenesis (Gouw et al., 2017) and FASN inhibition is detrimental to several cancer types (Baenke et al., 2013; Currie et al., 2013; Rohrig and Schulze, 2016). We reported that acyl-CoA synthetase long chain family member 3 (ACSL3), which synthesizes fatty acyl-CoA esters, is required for the viability of KMLC both *in vitro* and *in vivo* (Padanad et al., 2016). Also, recent studies indicated that lipid metabolism mediates adaptations to oncogenic *KRAS^G12C^* inhibition (Santana-Codina et al., 2020). However, the mechanisms governing the interaction of KM with FA metabolism and their functional consequences have not been characterized in sufficient detail to inform cancer therapy.

FA are required for the synthesis of complex lipids, such as phospholipids (PL) or triacylglycerides (TAG), which are used to build cellular membranes, to produce ATP through ß*-* oxidation (Baenke et al., 2013), or are stored in lipid droplets (Carracedo et al., 2013; Currie et al., 2013). On the other hand, in the presence of reactive oxygen species (ROS), polyunsaturated FA (PUFA), long-chain FA with multiple double bonds, undergo lipid peroxidation causing ferroptosis, a form of nonapoptotic ROS-dependent programmed cell death (Yang et al., 2016). Ferroptosis plays an important role in the regulation of cell survival in several physiologic as well as pathologic processes, including cancer (Zheng and Conrad, 2020).

Three hallmark features define cancer cell sensitivity to ferroptosis: the presence of oxidizable PL containing PUFA (PUFA-PLs), redox-active iron and inefficient lipid peroxide repair (*e.g.* glutathione peroxidase 4, GPX4) (Dixon and Stockwell, 2019; Liu et al., 2018; Yang et al., 2014). Consequently, the genes and metabolic pathways controlling lipid peroxide repair, antioxidant response, or PUFA metabolism mediate the sensitivity to ferroptosis.

The Lands cycle is the main process through which PL are remodeled to modify their FA composition (Lands, 1960). This process is conserved in the plant and animal kingdoms allowing the generation of new PL, effectively bypassing *de novo* synthesis of the entire PL molecule. Importantly, lung tissue synthesizes dipalmitoyl-PC (the major component of pulmonary surfactant) through this pathway, by de-acylating PC at the sn_2_ position and substituting a palmitic acid (C16:0) moiety at this site. Moreover, the Lands cycle constitutes the major route for incorporation and release of free arachidonic acid (AA, C20:4) and other PUFA into cellular PL, a process that is dependent on intracellular phospholipase A2 (PLA2). Hence, the proper regulation of the Lands cycle is important to control the accumulation of potentially toxic PL and FA, in order to maintain the integrity of cellular membranes (Ferrara et al., 2019; Wang and Tontonoz, 2019).

Here, we demonstrated that KMLC has high levels of PL decorated with saturated FA (SFA) and monounsaturated FA (MUFA). Moreover, FASN inhibition drastically reduces SFA/MUFA availability, obligating KMLC cells to incorporate the highly reactive PUFA into PL, thus decreasing the threshold for ferroptosis. Notably, silencing the key regulators of the Lands cycle phospholipase A2 group IVC (*PLA2G4C*) and lysophosphatidylcholine acyltransferase 3 (*LPCAT3*), induces ferroptosis in KMLC cells. We demonstrated that these processes are KM-dependent and confirmed our conclusions with metabolic flux analysis. This study provides the rationale for targeting FA synthesis and the Lands cycle as an effective therapeutic strategy for KMLC.

## Materials and Methods

### Human LC samples and Human LC PDXs

Human LC samples (n=6, Supplementary Table S1) were obtained from University of Cincinnati Biorepository under usage agreement (SR ID:TB0109). Frozen human LC PDXs (n=11, Supplementary Table S1) were obtained from Hamon Cancer Center at UT Southwestern Medical Center. Representative CT-scan images of a patient enrolled in the NCT03808558 trial were kindly provided by Dr. Gerber.

### Cell lines

LC cell lines were obtained from the cell line repository of the Hamon Center for Therapeutic Oncology Research (UT Southwestern Medical Center (*1*,*2*), IMR-90 human lung fibroblasts were from ATCC, Dr. Monte M. Winslow kindly provided mouse KMLC cell lines 238N1, 802T4, 368T1, 593T5 derived from LSL-*Kras^G12D^* lung tumors (Winslow et al., 2011). Cells were maintained as previously described (Konstantinidou et al., 2013; Padanad et al., 2016). For drug treatments, growth medium was replaced by HyClone RPMI medium (GE, SH30027) supplemented with 5% heat-inactivated dialyzed FBS (SIGMA-ALDRICH, F0394).

### Drugs and enzymatic inhibitors

TVB-3664 (FASNi) or the inactive isomer (TVB-2632) were from Sagimet Biosciences (San Mateo, CA). ML162 (#20455) or Ferrostatin-1 (#17729) were from Cayman. ARS-1620 (HY-U00418) was from MedChemExpress. 16:0-18:1 PC (hereafter PC #850457P), Sodium palmitate (#P9767) and N-Acetyl-L-cysteine (#A7250) were from Millipore-Sigma. CellROX™ Green Reagent (#C10444) was from Thermo Fischer Scientific.

### Mouse studies

All animal studies were approved by the Institutional Animal Care and Use Committee (IACUC) at University of Cincinnati (protocol 18-04-16-01). Both male and female mice were included in the analysis. Power calculation was conducted using ClinCalc.com using mean tumor burden from previous mouse experiments (Konstantinidou et al., 2009; Padanad et al., 2016). The investigators were blinded during post-study data analyses.

### Transgenic mice

*CCSP-rtTA/Tet-op-Kras^G12D^* (FVB/SV129 mixed background) mice were described previously (Fisher et al., 2001). We obtained lung-specific *Kras^G12D^* expression (i.e. KM) by feeding mice with doxycycline (doxy) -implemented food pellets (ENVIGO, TD 2018, 625 Dox, G) for 2 months. At this point, mice were randomly assigned to vehicle or FASNi. Lungs of *CCSP-rtTA/Tet-O-EGFR^L858R^* mice were kindly provided by Dr. Katerina Politi.

### Xenograft mouse models

We performed xenograft studies in 6-week-old NOD/SCID mice (Jackson Laboratory), as previously described (Konstantinidou et al., 2013; Padanad et al., 2016). Briefly, 1 × 10^6^ cells (A549 or H460) were injected in the right flank of NOD-SCID mice. Tumors were measured twice a week using a digital caliper. Mice were euthanized when xenografts reached 2 cm^3^.

### FASNi in vivo treatment

TVB-3664 (60 mg/kg/100 μL/mouse) was dissolved in 0.2% DMSO-PBS, sonicated and administered daily via oral gavage. We treated *CCSP-rtTA/Tet-op-Kras^G12D^* mice after feeding them with doxy for 2 months. We treated NOD/SCID mice when xenografts reached ∼30-50 mm^3^. At this time, mice were randomly assigned to either vehicle (0.2% DMSO in PBS, control) or FASNi (TVB-3664) group.

### Blood and tissue collection

We collected blood (150 µL) from the submandibular vein and left at RT until clot formation. Lung were harvested as previously described (Fisher et al., 2001). Briefly, we cannulated the right heart and perfused anesthetized with PBS lacking calcium and magnesium. Lung lobes were excised, then either fixed overnight in 4% paraformaldehyde at 4°C, or snap frozen and stored at −80°C for further analyses.

### Viability assays

For MTT assays, we plated 5000 cells/0.1 mL/well in 96-well plates. The day after, we added 0.1 mL of medium containing appropriate concentrations of the compound under study to each well. After 7 days (or 5 days for ARS-1620) of treatment, 10 μL of MTT solution (5 mg/mL) was added into each well. After 4 hrs incubation, the medium was removed and the formazan salts dissolved in 200 μL DMSO/ well for 10 min at 37°C. The absorbance was read at 570 nm (OD 570 nm). Data are represented as mean ± SD of 3 independent experiments.

*Crystal violet assays* were performed as previously described (Feoktistova et al., 2016). Briefly, 3-4 x 10^4^ cells/ well were seeded in 12-well plate. At the experimental endpoint, 0.25 mL/well of crystal violet solution (0.5% of crystal violet powder in 20% methanol) was added to the plate and incubated at RT for 20 min on an orbital shaker. After removing the crystal violet solution, the plate was washed with water and let to dry overnight. Pictures were taken before adding 1 mL/well of methanol. After shaking 20 min at RT, the absorbance was read at 570 nm (OD 570 nm).

### Cell cycle analysis

We performed cell cycle analysis after 4 days of pharmacological treatments, or 72 hrs after siRNA transfection or doxy-dependent induction of shRNAs, respectively. The cells were fixed in ice-cold 70% ethanol for 30 min. RNA digestion was performed with 100 g/mL RNase A (Sigma Aldrich, R6513) for 15 min at 37°C. DNA staining was performed with 50 μg/mL propidium iodide (Sigma Aldrich, P4170) for 30 min at 37°C. The cells were analyzed using a BD LSRFortessa™ Flow Cytometer, and the cell cycle distribution was determined with FlowJo v8.7 Software.

### Oil Red Oil Staining

We treated cells either with FASNi, 0.2 µM or with DMSO, 0.2% (i.e. vehicle) for 4 days. Cells were rinsed with 1X PBS twice, fixed with formalin (3%) for 1 hour, and washed again. Cells were incubated with isopropanol (60%) for 5 minutes and then with ORO solution (ORO, 2mg/mL, Alfa Aesar, A12989) for 20 min. Nuclei were stained with hematoxylin stain solution, Gill 1(RICCA, #3535-16). For quantification, ORO staining was extracted with 100% isopropanol. The absorbance was read at 492 nm.

### Protein extraction and Immunoblot

Tissues or cell lines were lysed in ice-cold RIPA lysis buffer containing cOmplete^TM^ protease inhibitors cocktail and PhosSTOP^TM^ phosphatase inhibitors cocktail (Millipore Sigma). 30 µg of total protein were separated using Midi Criterion TGX Stain-Free precast gels (Bio-Rad, 5678024), transferred onto nitrocellulose membranes (Bio-Rad, 1620112), and then blocked with 5% Blotting-Grade Blocker (Bio-Rad, 1606404) in TBS for 1 h, at RT. The membranes were incubated overnight at 4°C with the indicated primary antibodies: FASN (C20G5) rabbit mAb (CST, 1:1000 dil); SCD1 (C12H5) rabbit mAb (CST, 1:1000 dil); AMPKα Antibody rabbit pAb (CST #2532, 1:1000 dil); phospho-AMPKα (Thr172) (40H9) rabbit mAb (CST, 1:1000 dil); ACC1 (C83B10) Rabbit mAb (CST, 1:1000 dil); Phospho-ACC1 (Ser79) (D7D11) rabbit mAb (CST, 1:1000 dil); KRAS (234-4.2) mouse mAb (Millipore Sigma, 1:700 dil); pan RAS (C-4) mAb (SCBT, 1:500 dil); Cofilin (D3F9) rabbit mAb (CST, 1:1000 dil). Cofilin was used as loading control. Either goat anti-rabbit or anti-mouse IgG (H+L) cross adsorbed, DyLight® 800 (Thermo Scientific, 1:5000 dil) was used as secondary antibody. Blots were analyzed using Odyssey Scanner and Image Studio Software (LI-COR).

### Malonyl-CoA quantification

Malonyl-CoA concentration was assessed using the human malonyl Coenzyme A ELISA kit (MyBiosource, MBS705079), following manufacturer’s instructions. Test samples were prepared in triplicates.

### Measurement of fatty acid oxidation (FAO)

We used a fatty acid oxidation colorimetric assay kit (Biomedical Research Service Center, State University of New York, E-141) as previously described (Kwong et al., 2019). Samples were assayed in triplicate.

### NADPH quantification

NADPH in lysates of cell lines or A549 xenografts was determined using a colorimetric NADPH Assay Kit (Abcam’s, ab186031), following the manufacturer’s instructions on LC cells and A549 xenografts treated with either FASNi or vehicle for 4 days. Test samples were prepared in triplicates.

### AMP quantification

We used a colorimetric AMP colorimetric assay kit (BioVision, K229-100) to detect AMP in cell lysates (∼1 x 10^7^) and A549 xenografts (∼10 mg) following the manufacturer’s instructions. Samples were assayed in triplicate. 1 mM AMP Standard was used as positive control.

### Cu(I)-catalyzed azide-alkyne cycloaddition reaction (Click-iT chemistry)

Cells were grown on coverslips in a 12-well plate and kept either with FASNi, 0.2 µM or with vehicle (DMSO, 0.2%) for 4 days. Then, we incubated them with 20 µM arachidonic acid alkyne (Cayman Chemical, 10538) in RPMI with 2% ultra-fatty acid free BSA (SIGMA ALDRICH, A6003-25G) for 6 hrs. After fixation with 3% paraformaldehyde in PBS for 15 min and permeabilization with 0.25% Triton X100 in PBS for 15 min, cells were washed with 1% BSA in PBS. The Click-iT reaction cocktail was prepared according to manufacturer’s instructions(Kolb and Sharpless, 2003). Briefly, Click-iT™ Cell Reaction Buffer Kit (Thermo Scientific, C10269) was mixed with CuSO4 (1 mM final concentration) and Alexa-Fluor 488 azide (10 µM final concentration, Thermo Scientifc, A10266). After staining nuclei with DAPI, cells were washed again, and mounted on a slide using Fluoromount-G medium.

### C11-BODIPY staining

We stained fixed cells with 2.5 µM C11-BODIPY581/591 (SIGMA ALDRICH, D-3861) for 60 min. The ROS-dependent oxidation of the polyunsaturated butadienyl portion of this lipid probe results in a shift of the fluorescence emission peak from ∼590 nm (reduced: red) to ∼510 nm (peroxided: green) (Drummen et al., 2002; Pap et al., 1999). Briefly, cells were treated with 0.2 µM FASNi or 0.2% DMSO, for 4 days. In rescue experiments, cells were co-treated with FASNi and one of the following: palmitate (100 µM), phosphatidylcholine (PC, 100 µM), or Ferrostatin-1 (Fer-1, 1 µM) for 4 days. 5 mM N-acyl-cysteine (NAC) was added for 60 min after 4 days of FASNi treatment. For snap-frozen samples, we used 10 µm-thick cryo-sections of A549 xenografts (5 mice/group) and lungs of CCSP-rtTA/Tet-op-Kras (3 mice/group). Samples were incubated with C11-BODIPY581/59, washed with PBS, fixed with 10% formalin for 1 hour, counterstained with DAPI and mounted using Fluoromount-G medium (Thermo Scientific). For cells, three images per slide were acquired using a Zeiss LSM 710 confocal microscope equipped with a Plan-Apo 63x/1.4 oil DIC M27 objective. For tissues, three images per sample were acquired using a Lionheart FX (BioTek) and Gen5 software (v3.06). We quantified the images with ImageJ software (version 1.46; NIH, Bethesda, MD, USA).

### Plasmids and virus production

*pBabe-Kras^G12D^* (#58902) and LT3GEPIR (#111177) were from Addgene. The insert expressing the shRNA was cloned into the vector *LT3GEPIR* in the XhoI and EcoRI sites. The shRNAs target sequences were selected either from the library described by Feldmann et al. (Fellmann et al., 2013) or from the splashRNA database (Pelossof et al., 2017) (http://splashrna.mskcc.org/). The hairpin targeting sequences are: *shLPCAT3 #1* 5’-GCCTCTCAATTGCTTATTTTA-3’; *shLPCAT3 #2* 5’-AAGGAAAGAGAAGTTAAA-3’; *shPLA2G4C #1* 5’-CAGAATGAATGTGATAGTTCA-3’; *shPLA2G4C #2* 5’-ACATGGTTATCTCTAAGCAAA-3’; *shGPX4 #1* 5’-GTGGATGAAGATCCAACCCAA-3’; *shGPX4 #2* 5’-AGGCAAGACCGAAGTAAACTA-3’. *shKRAS #10* 5’-AAGTTGAGACCTTCTTAATTGGT-3’; *shKRAS #40* 5’-TCAGGACTTAGCAAGAAGTTA-3’. Production of lentiviruses was performed as previously described (Padanad et al., 2016).

### siRNA

Predesigned FASN siRNAs were from Millipore Sigma (#1, SASI_Hs01_00057850; #2, SASI_Hs01_00057851; #3, SASI_Hs02_00336920; #4, SASI_Hs01_00057849). A custom siRNA library (Supplementary Table S2), universal MISSION® siRNA Universal Negative Control #1 (cat. SIC001) and #2 (cat. SIC002) were from Millipore Sigma. We used the DharmaFECT 4 Transfection Reagent (Thermo Scientific) for siRNA transfection. RNA was extracted using TRIzol Reagent (cat. 15596, Life Technologies) and retrotranscribed using iScript cDNA Synthesis kit (cat. 170-8891, Biorad). Knock-down efficiency was evaluated 48 hrs after transfection via real-time PCR using the PowerUP^TM^ SYBR® Green Master Mix (cat. A25742, Thermo Fisher Scientific) and custom designed primers (Supplementary Table S2).

### RNA-seq and bioinformatic analysis

We extracted total RNA with TRIzol. We removed residual genomic DNA was removed with the Turbo DNA-free kit (AM1907, ThermoFisher). One µg of total DNase-treated RNA was used for library preparation with the NEBNext Ultra II Directional RNA Library Prep kit. Reads were aligned to the human hg38 reference genome using STAR (v3.7.3a) (Dobin et al., 2013). Gencode annotation for human (version v37) was used as reference alignment annotation and downstream quantification. Gene level expression was calculated using featureCounts (v2.0.1) (Liao et al., 2014) using intersection-strict mode by exon. Counts were calculated based on protein-coding genes from the annotation file. Low expressed genes were filtered using a per time-point approach with RPKM >= 0.5 in all samples in one or the other time-point. Differential expression was performed in R using *DESeq2* (Love et al., 2014). Surrogates variables were calculated using *sva* (Leek et al., 2012) and included in the modelling. We estimated log2 fold changes and P-values. P-values were adjusted for multiple comparisons using a Benjamini-Hochberg correction (FDR). Differentially expressed genes were considered for FDR < 0.05. Gene list enrichment analysis was carried out using Enrichr(Chen et al., 2013; Kuleshov et al., 2016) (https://amp.pharm.mssm.edu/Enrichr/<u>#</u>).

### Laser-capture microdissection (LMD) of tumor tissue

LMD was performed at the Histopathology Core of UT Southwestern Medical Center as previously described (Bonner et al., 1997). Briefly, 10 µm cryo-sections of tumors were mounted onto PEN membrane glass slides (Thermo Fisher Scientific, LCM05220Sections were then stained using Histogene™ Staining Solution (Thermo Fisher Scientific, KIT0415), washed with HPLC-grade water and subjected to LMD using a Leica LMD System. LMD sections were recovered in 0.2 mL tubes, immediately snap-frozen and stored at −80°C until the analysis. 4 replicates per sample were prepared.

### Solvents and reagents

All the HPLC or LC/MS grade solvents were from Sigma-Aldrich (St Louis, MO, USA). SPLASH LipidoMix™ standards, were from Avanti Polar Lipids (Alabaster, AL, USA). Fatty acid (FA) standards (FA(16:0{^2^H_31_}), FA(18:1ω9{^2^H_5_}) and FA(20:4 ω6{^2^H_8_}) were from Cayman Chemical (Ann Arbor, MI, USA). An eVol® precision pipette equipped with a glass syringe (Trajan Scientific, Austin TX, USA) was used for the addition of FA standards.

### Sample preparation for MS/MS^ALL^ and GC/MS

Laser-captured microdissected samples, cell pellets containing 2.5 ×10^5^ cells or 10 µL of serum were transferred to glass tubes for Liquid-Liquid Lipid Extractions (LLE). For xenografts, 100 mg of tissue were transferred to a 2.0 mL pre-filled Bead Ruptor tube (2.8 mm ceramic beads, Omni International, Kennesaw, GA, USA), and homogenized in 1mL of methanol/dichloromethane (1:2, v/v) using a Bead Ruptor (Omni International). Aliquots equivalent to 0.5 mg of tissue were used for LLE.

### MALDI Imaging Mass Spectrometry (MALDI-IMS)

10-µm-thick sections were mounted on PEN membrane glass slides and stored at −80°C. They were processed by the Chemical Imaging Research Core, UT MD Anderson Cancer Center. Matrix was applied using the sublimation method with a Shimadzu IM Layer (Shimadzu North America, Columbia, MD). Sampling was performed using DHB for positive mode and 9-AA for negative mode. We used 300 laser shots at 1 KHz using a laser pulse energy with an average of 25 μJ. For DHB and 9-AA, the laser was adjusted to 60% and 50%, respectively. The mass range was 50-1200 m/z, and the instrument was calibrated using peak signals from red phosphorus. For MS/MS MSI analysis of taurodeoxycholic acid, parent ion atm/z 498.295 was selected, and fragment ion at m/z 124.006 was monitored at a collision trap voltage of 40 V. The laser shots were 500 shots per spot at 50% power. Data were acquired using a Waters Synapt G2 Si (Waters Corporation, Milford, MA) at 60-μm spot size with 100-μm spacing. For relative quantitation, the MSI data were converted to msIQuant format via imzML. Regions of interest (ROI) were annotated manually in msIQuant version 2.0 (Källback et al., 2016). Total ion current (TIC) normalized average intensities of metabolites were exported from each region for statistical analysis. Peak identification was manually done using LIPID MAPS Lipidomic Database (LMSD).

### Lipidomic experiments

After 4 days of treatment, with vehicle or FASNi, cells were either washed twice with cold PBS and harvested for MS/MS^ALL^, or incubated with 3 mM Ethyl Acetate-1,2 ^13^C_2_ (SIGMA ALDRICH, 283819) for 7 hours (1 hour for the time-lapse experiment) to measure *de novo* palmitate synthesis by GC/MS.

### Lipid profiling by direct-infusion MS/MS^ALL^

The LLE was performed at RT through a modified Bligh/Dyer extraction technique. Briefly, 3mL of methanol/dichloromethane/water (1:1:1, v/v) were added to the samples. The mixture was vortexed and centrifuged at 2671 g for 5 min. The organic phase (bottom phase) was collected and dried under N_2_. The extracts were resuspended in 600 µL of dichloromethane/methanol/isopropanol (2:1:1, v/v/v) containing 8mM ammonium fluoride (NH4F) and 33 µL of 3:50 diluted SPLASH LipidoMix™ internal standard. Extracts were infused into a SCIEX quadrupole time-of-flight (QTOF) TripleTOF 6600+ mass spectrometer (Framingham, MA, USA) via a custom configured LEAP InfusePAL HTS-xt autosampler (Morrisville, NC, USA). Electrospray ionization (ESI) source parameters were, GS1 25, GS2 55, curtain gas (Cur) 20, source temperature 300 °C and ion spray voltage 5500V and −4500V in positive and negative ionization mode, respectively. GS1 and 2 were zero-grade air, while Cur and CAD gas was nitrogen. Optimal declustering potential and collision energy settings were 120V and 40eV for positive ionization mode and −90V and −50eV for negative ionization mode. Samples were infused for 3 min at a flow rate of 10µL/min. MS/MS^ALL^ analysis was performed by collecting product-ion spectra at each unit mass from 200-1200 Da. Analyst® TF 1.7.1 software (SCIEX) was used for TOF MS and MS/MS^ALL^ data acquisition. Data analysis was performed using an in-house script, LipPy. This script provides instrument quality control information, isotopic peak corrections, lipid species identification, data normalization, and basic statistics.

### Fatty acid profiling by GC-MS

Total fatty acid profiles were generated by a modified GC-MS method previously described (Quehenberger et al., 2011). The lipid extract was spiked with 100uL of 0.5µg/mL FA standard mixture (FA(16:0{^2^H_31_}), FA(20:4 ω6{^2^H_8_}) and FA(22:6 ω3{^2^H_5_}) in methanol, then hydrolyzed in 1 mL of 0.5M potassium hydroxide solution prepared in methanol at 80 °C for one hour. Hydrolyzed FA were extracted by adding 2mL of dichloromethane/water (1:1, v/v) to the sample in hydrolysis solution. The mixture was vortexed and centrifuged at 2671 g for 5 min. The organic phase (bottom phase) was collected and dried under N_2_. For the free FA profile, the lipids were extracted by adding 500 µL of water, 500 µL of methanol containing 50 mM of HCL and 1mL of iso-octane to the glass tube containing the sample. The solution was spiked with 100 µL of 0.5 µg/mL fatty acid standard mixture, shacked for 5 min, centrifuged at 2671 g for 5 min, and the organic phase (upper phase) was collected to a fresh glass tube. The extraction procedure was repeated two times by adding 1mL of iso-octane to the mixture. The organic phases were pooled together and dried under N_2_. To analyze FA present in the polar lipid fraction, a three-phase extraction method was performed, as already described (Vale et al., 2019). Briefly, lipids were extracted with water, methyl acetate, acetonitrile (ACN) and Hexane (1:1:0.75:1). After centrifugation, polar lipids (upper phase) were collected and dried. Total or polar FA samples were resuspended in 50 µL of 1 % triethylamine in acetone, and derivatized with 50 µL of 1% pentafluorobenzyl bromide (PFBBr) in acetone at RT for 25 min in capped glass tubes. Solvents were dried under N_2_, and samples were resuspended in 500 µL of isooctane. Samples were analyzed using an Agilent 7890/5975C (Santa Clara, CA, USA) by electron capture negative ionization (ECNI) equipped with a DB-5MS column (40m х 0.180mm with 0.18µm film thickness) from Agilent. Hydrogen (carrier gas) flow rate was 1.6mL/min and injection port temperature was set at 300 °C. Sample injection volume was 1µL. Initial oven temperature was set at 150 °C, and then increased to 200 °C at a 25 °C/min, followed by an increase of 8 °C/min until a temperature of 300 °C was reached and held for 2.2 min, for a total run time was 16.7 min. FA were analyzed in selected ion monitoring (SIM) mode. The FA data were normalized to the internal standards. Fatty acid with carbon length C ≤ 18 were normalized to FA(16:0{^2^H_31_}), C = 20 were normalized to FA(20:4 ω6{^2^H_8_}), and C = 22 were normalized to FA(22:6 ω3{^2^H_5_}). Data were processed using MassHunter software (Agilent).

### HPLC-MS/MS

PDXs and human lung cancer specimens were stored at −80 °C. LLE was performed using a modified Bligh/Dyer extraction technique. The extracts were resuspended in 30 µL of dichloromethane/methanol/isopropanol (IPA) (2:1:1, v/v/v) containing 8mM NH4F and SPLASH LipidoMix™ internal standards. Reversed-phase chromatographic separation was achieved with the Acclaim C30 column: 3 μm, 2.1 × 150 mm (Thermo Fisher Scientific, Waltham, MA). The column was maintained at 35°C and tray at 20°C. Solvent A was composed of 10 mM ammonium formate (AF, LC-MS grade) in 60:40 Acetonitrile (ACN):water (LC-MS grade) with 0.1% formic acid (FA, LC-MS grade). Solvent B was composed of 10 mM AF with 90:10 IPA:ACN with 0.1% FA. The flow rate was 250 μL/min, and the injection volume was 10 μL. The gradient was 50% solvent A (3 to 50%). The Orbitrap (Thermo) mass spectrometer was operated under heated electrospray ionization (HESI) in positive and negative modes separately for each sample. The spray voltage was 3.5 and 2.4 kV for positive and negative mode, the heated capillary was held at 350 °C and heater at 275°C. The S-lens radio frequency (RF) level was 45. The sheath gas flow rate was 45 units, and auxiliary gas was 8 units. Full scan (m/z 250–1200) used resolution 30,000 at m/z 200 with automatic gain control (AGC) target of 2 × 105 ions and maximum ion injection time (IT) of 100 ms. Normalized collision energy (NCE) settings were 25, 30, 35 %.

Lipid identification and relative quantification were performed with LipidSearch 4.1 software (Thermo) as previously described (Breitkopf et al., 2017). The search criteria were as follows: product search; parent m/z tolerance 5 ppm; product m/z tolerance 10 ppm; product ion intensity threshold 1%; filters: top rank, main isomer peak, FA priority; quantification: m/z tolerance 5 ppm, retention time tolerance 1 min. The following adducts were allowed in positive mode: +H, +NH4, +H____H2O, +H____2H2O, +2H, and negative mode: ____H, +HCOO, +CH3COO, -2H.

### Statistical analysis

Data analysis was performed using Microsoft Excel for Mac (version 16.34) and GraphPad Prism version 9 (GraphPad Software, San Diego, CA, USA, www.graphpad.com). All data presented are expressed as mean ± SEM or ±SD of three or more biological replicates/group (n values in each figure/figure legend). The significance of the results was assessed using two-tailed unpaired Student’s t test to compare two groups. When more than two groups were compared, one- or two-way ANOVA was used followed by Dunnett’s, Tukey’s or Sidak’s post-test.

## Results

### Mutant KRAS induces a specific lipid profile in lung cancer

*CCSP-rtTA/Tet-O-Kras^G12D^* (hereafter *TetO-Kras^G12D^*) and *CCSP-rtTA/Tet-O-EGFR^L858R^* (hereafter *TetO-EGFR^L858R^*) mice, when fed doxy, invariably develop lung tumors which recapitulate tumorigenesis and histological features of human LC (Fisher et al., 2001; Politi et al., 2006). We performed mass spectrometry (MS) analysis on micro-dissected lung tumors and unaffected parenchyma (Fig. 1A). We found that the two tumor types have different lipidomic signatures. *TetO-Kras^G12D^* LC shows a significant increase in PC and TAG, as well as of sphingomyelins (SM) and phosphatidylethanolamine (PE), and a decrease in lysophosphatylcholines (LysoPC), while *TetO-EGFR^L858R^* preferentially has high phosphatidylinositol (PI) and low TAG, (Fig. 1B, Supplementary Fig. S1A). Nevertheless, both *TetO-Kras^G12D^* and *TetO-EGFR^L858R^* tumors have higher cholesteryl-esters (CE) and lower diacylglycerides (DAG) then healthy lung (Fig. 1B, Supplementary Fig. S1A). In particular, *TetO-Kras^G12D^* tumors are enriched in PC species with SFA and MUFA acyl chains, but have less PUFA-containing PC (and PE) than healthy lung (Fig.1C). To determine the spatial distribution of the major lipid species identified by MS, we used Matrix Assisted Laser Desorption/Ionization (MALDI) imaging. We confirmed that *TetO-Kras^G12D^* tumors are enriched in TAG, SM and PC (Fig. 1D). In particular, we observed that SFA- and MUFA-PC are preferentially localized within the tumors, while PUFA-PC accumulate in the surrounding lung parenchyma (Fig. 1D). Notably, we observed the same lipidomic pattern in KMLC patient-derived xenografts (PDXs) and primary patient specimens (Fig. 1E-G; Supplementary Fig. S1B).

**Figure 1.**
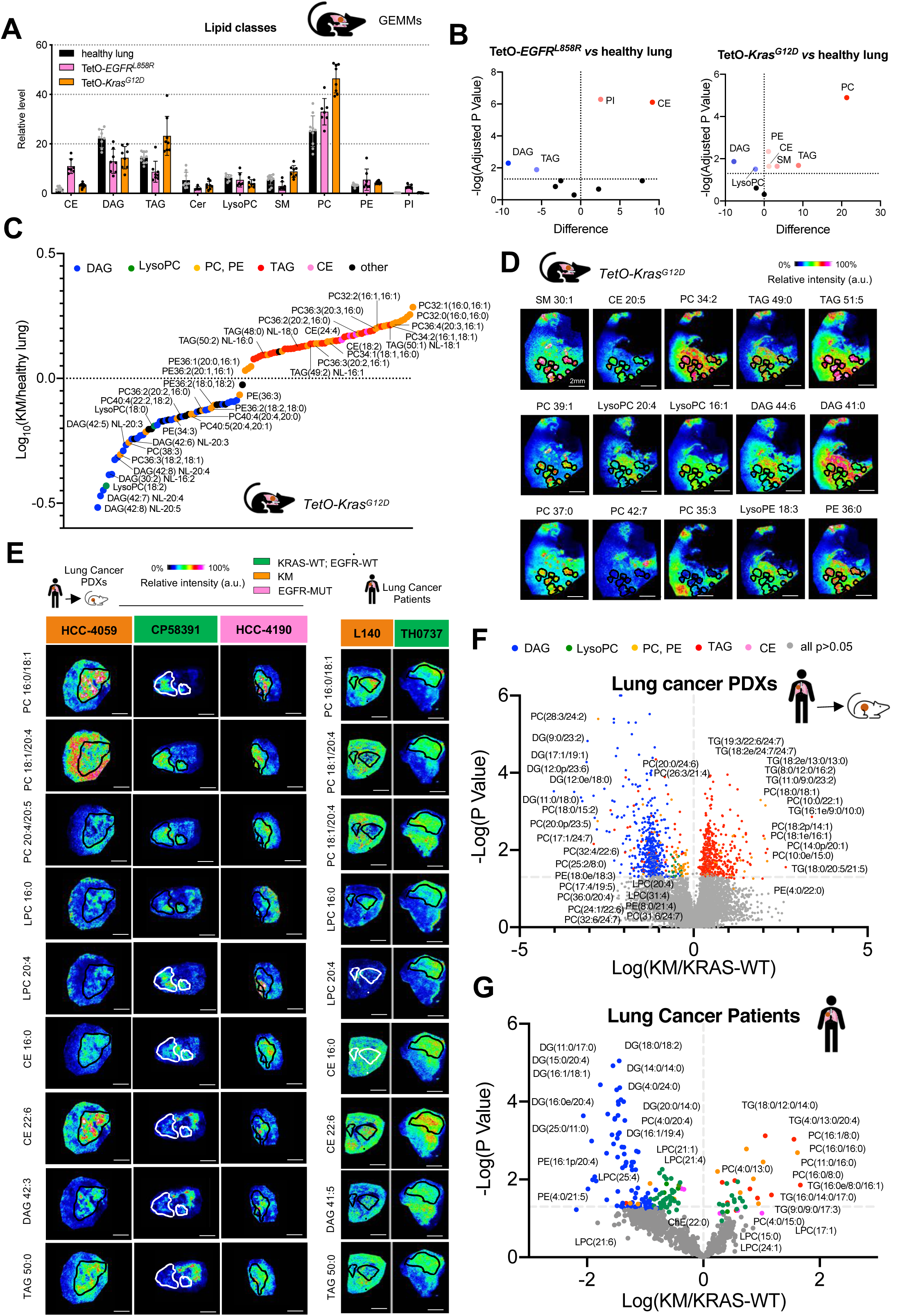
KMLC has a specific lipidome in mouse and human samples. **(A, B)** Lipidomic analysis of murine TetO-*Kras^G12D^* tumors (n=4), TetO-*EGFR^L858R^* (n=3) and unaffected healthy lung (n=3). Each dot indicates a lung/tumor section. Data are expressed as mean ± SD. PC, phosphatidylcholines; TAG, triglycerides; PE, phosphatidylethanolamines; CE, cholesteryl-esters; PI, phosphatidylinositols; DAG, diacylglycerides; SM, sphingomyelins; Cer, ceramides; LysoPC, lysophosphatidylcholines. Volcano plots in (B) show the lipid classes that are differentially represented for each comparison. The adjusted P value and difference were calculated using multiple t tests with alpha =0.05. **(C)** MS/MS and **(D)** MALDI imaging analyses showing lipid differentially represented in TetO-*Kras^G12D^* tumors as compared to unaffected heathy lung. **(E)** Representative pictures of MALDI imaging analysis of lung cancer patient-derived xenografts (PDXs) and primary human lung cancer specimens of the indicated genotype. **(F, G)** HLPC-MS/MS analysis of lung cancer PDXs and primary human lung cancer specimens of the indicated genotype. Volcano plots show lipid species identified by HLPC-MS/MS differentially represented in KM *versus* KRAS-WT samples (PDXs, KM n=5 and KRAS-WT n=4; lung cancer patients, n=3/group). P values and difference were calculated using multiple t tests (p<0.05). In (D) and (E) rainbow scale represents ion intensity normalized against the total ion count (%) and black circles represent the major tumor areas.

These data suggest that KM increases the intratumor availability SFA and MUFA for the synthesis of TAG and PC (Supplementary Fig. S1C).

### Mutant KRAS induces a dependency on *de novo* lipogenesis

To test whether the lipidome of KMLC depends on *de novo* lipogenesis, we used TVB-3664 (Sagimet Biosciences), a specific FASN inhibitor (FASNi), and *FASN* silencing on a panel of human and mouse LC-derived cell lines (Gazdar et al., 2010; Phelps et al., 1996; Winslow et al., 2011) (Fig. 2A and Supplementary Fig. S2). Both approaches caused a G2/M cell cycle arrest in KMLC, without affecting KRAS-WT and EGFR-MUT LC cells (Fig. 2B; Supplementary Fig. S2). Notably, exogenous palmitate rescues the detrimental effects of FASNi, while an inactive FASNi isomer does not affect the viability of KMLC cells (Fig. 2A and 2B). As previously reported (Gouw et al., 2017), we found that KM correlates with FASN overexpression in *TetO*-*Kras^G12D^* mice (Supplementary Fig. S3A), in human KMLC cell lines (Supplementary Fig. S3B) and two independent tumor microarrays (Supplementary Table S3; Supplementary Fig. S3C-S3F), while there is no significant correlation with EGFR-MUT (Supplementary Fig. S3G and S3H).

**Figure 2.**
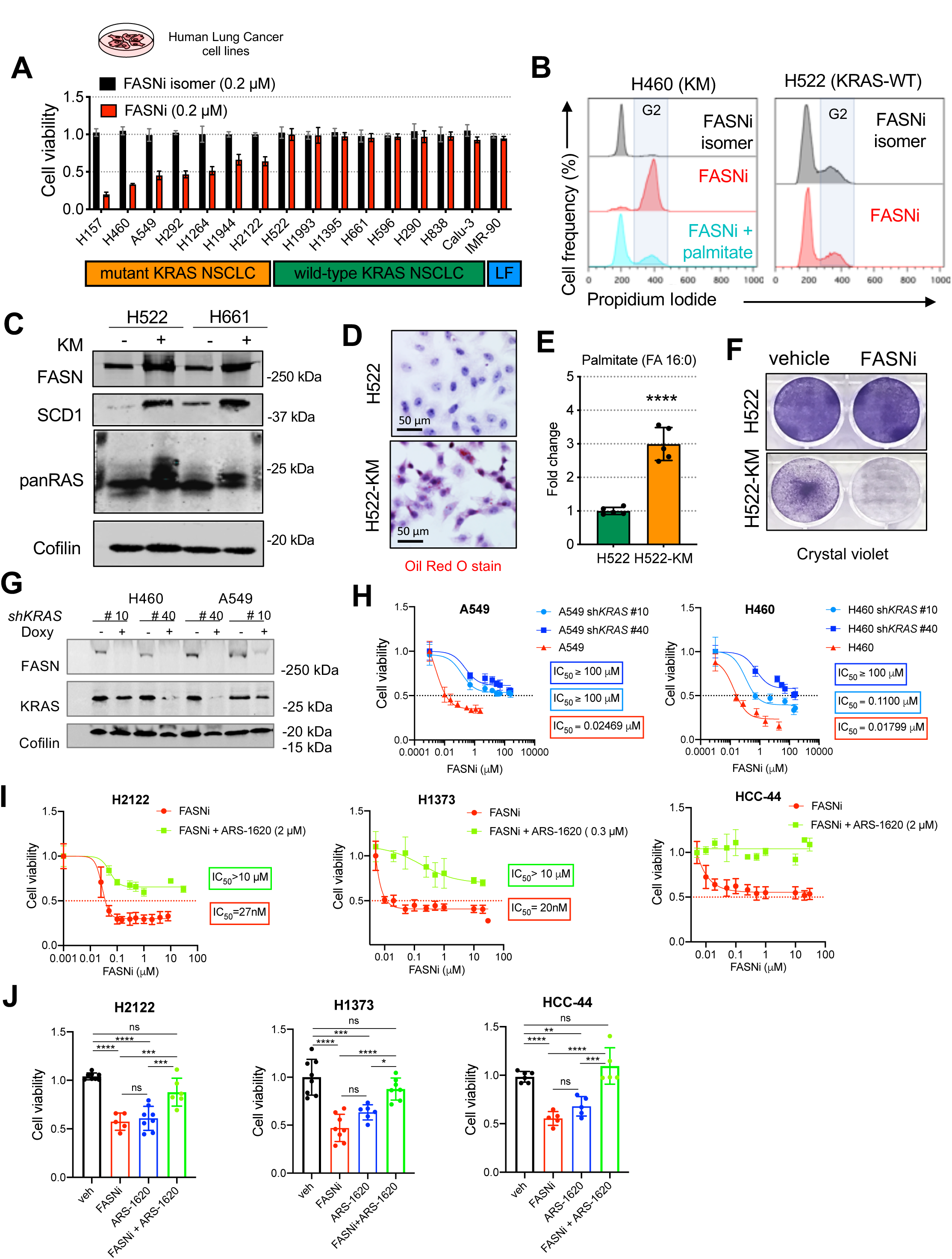
KM is required to induce dependency on FASN. **(A)** Viability assay of human LC derived cell lines. Cell line, genotype and treatments are indicated. LF: lung fibroblasts. **(B)** Cell cycle analysis of H460 and H522 cells, as representative examples of KM and KRAS-WT LC cells, treated as indicated. **(C)** Immunoblot of FASN, SCD1 and pan-RAS in H522 and H661 cells transduced as indicated. (**D**) Oil red O staining, relative steady-state quantification of palmitate (FA 16:0) **(E)** and crystal violet assay **(F)** of H522 and H522-KM cells treated as indicated. (**G**) Immunoblot of FASN and KRAS in H460 and A549 cells transduced with doxy-inducible shRNAs targeting *KRAS*. **(H**) MTT viability assay of H460 and A549 cells treated with FASNi before and after induction of *KRAS* knock-down. **(I**) MTT viability assay of H2122, H1373 and HCC-44 cells (*KRAS^G12C^* mutant) treated with FASNi alone or in combination with the *KRAS^G12C^* inhibitor ARS-1620. Data are expressed as mean ± SD. In (E) Student t test with ****p<0.0001. In (J) one-way ANOVA followed by Tukey’s multiple comparison test with ***p<0.001 and ****p<0.0001.

To test whether *KM* expression is sufficient to establish a dependency on FASN, we ectopically expressed *KM* in H522 and H661 KRAS-WT LC cells. KM induces upregulation of FASN and its downstream enzyme, stearoyl-CoA desaturase 1 (SCD1) (Fig. 2C), accumulation of lipid droplets (Fig. 2D; Supplementary Fig. S4A) and palmitate (Fig. 2E), sensitivity to FASN inhibition and *FASN* silencing (Fig. 2F; Supplementary Fig. S4B and S4C). On the contrary, *KRAS* silencing induces a significant decrease of FASN in KMLC cells (Fig. 2G), which also become resistant to FASNi (Fig. 2H). Furthermore, the *KRAS^G12C^* inhibitor ARS-1620 reverses the effects FASNi in *KRAS^G12C^* LC cell lines (Fig. 2I and 2J). All together these data demonstrate that KM causes LC cells to become dependent on FASN.

### FASNi inhibits *de novo* lipogenesis independently of KRAS mutational status

To gather mechanistic insights into the differential sensitivity to FASNi, we measured the drug uptake, stability and its activity in LC cells.

Irrespective of the *KRAS* mutational status, FASNi readily accumulates intracellularly (Supplementary Fig. S5A and S5B), inhibits *de novo* FA synthesis (Fig. 3A; Supplementary Fig. S5C), causes a concomitant accumulation of malonyl-CoA (FASN substrate) and NADPH (FASN cofactor) (Fig. 3B and 3C), depletion of lipid droplets (Fig. 3G; Supplementary Fig. S5D) and downregulation of ß-oxidation (Fig. 3E). Moreover, FASNi causes accumulation of AMP (Fig. 3F), triggering the phosphorylation of AMPK and ACC1 in both genotypes (pACC1^S79^) (Fig. 3G). This observation is consistent with the notion that AMPK limits FA synthesis through direct phosphorylation of ACC1, the enzyme that catalyzes the synthesis of malonyl-CoA, the substrate of FASN (Garcia and Shaw, 2017). Thus, the activation of AMPK/ACC1 enhances the effect of FASNi on FA synthesis (Fig. 3H). However, since these effects occur in both KM and KRAS-WT LC cells, they do not provide an explanation for the specific dependency of KMLC cells on FASN.

**Figure 3.**
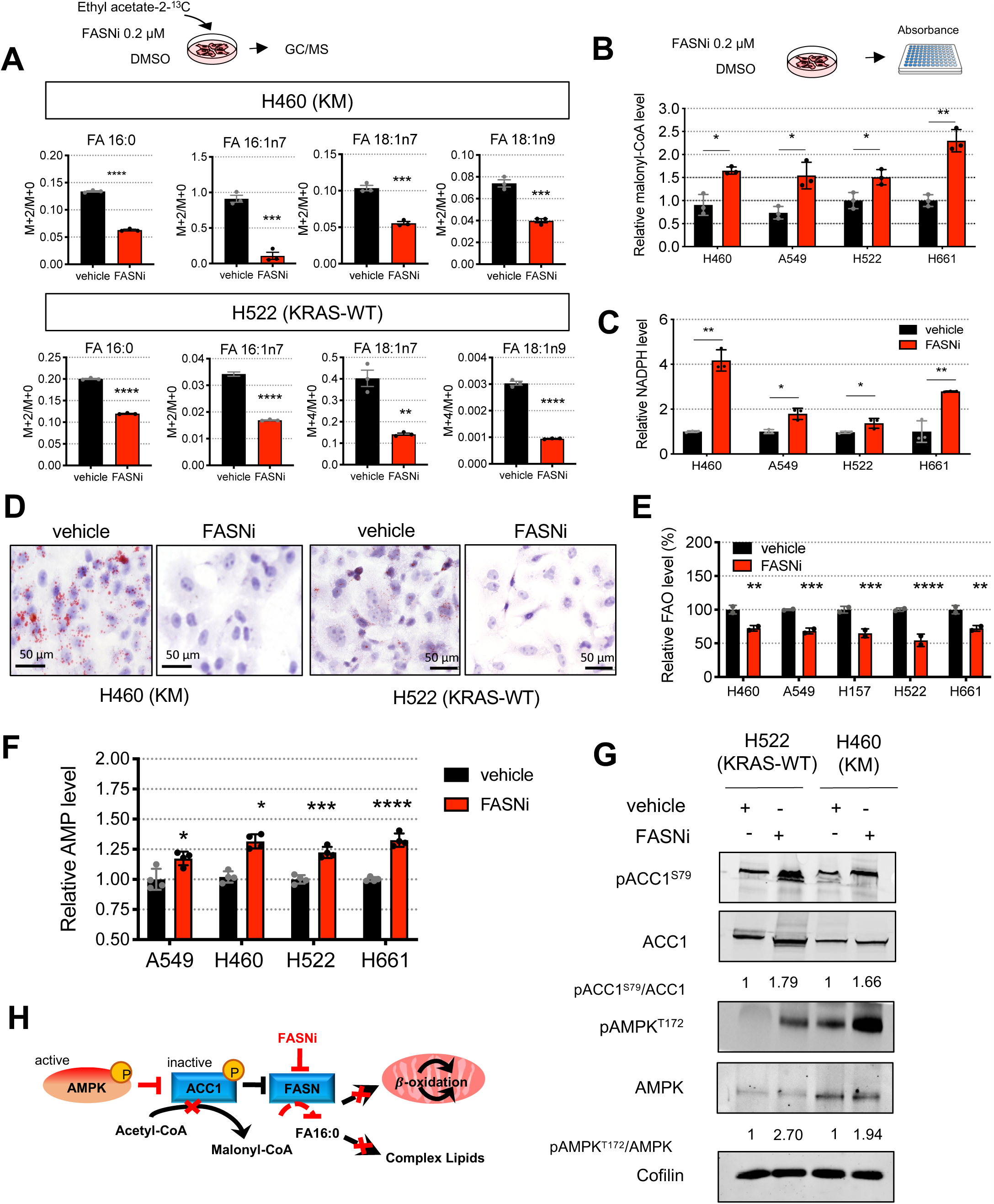
FASNi inhibits fatty acid synthesis and ß-oxidation in both KM and KRAS-WT cells. **(A)** GC/MS quantification of newly synthesized FA in H460 and H522 cells after overnight ethyl acetate-2-^13^C labelling, treated as indicated. Either M+2/M+0 or M+4/M+0 ratio is reported. Palmitate, FA 16:0; palmitoleate, FA 16:1n7; vaccenate, FA 18:1n7; oleate, FA 18:1n9 (n=3 independent experiments). **(B, C)** Relative quantification of malonyl-CoA and NAPH of vehicle- and FASNi-treated LC cells (n=3 independent experiments). **(D)** Oil red O staining for lipid droplets in H460 and H522 cells. **(E, F)** Relative quantification of FA ß-oxidation (FAO) and AMP in the indicated cells treated with vehicle or FASNi. **(G)** Immunoblot of phospho-Ser79-ACC1 (pACC1^S79^), ACC1, FASN, phosphor-Thr172-AMPK (pAMPK^T172^) and AMPK. **(H)** Schematic of the AMPK/ACC1/FASN axis. FASNi inhibits the synthesis of palmitate (FA 16:0) thereby blocking the synthesis of complex lipids and ß-oxidation (FAO). These events trigger the activation of AMPK which in turn phosphorylates and deactivates ACC1. ACC1 phosphorylation blocks the synthesis of malonyl-CoA (the substrate for FA 16:0 synthesis) potentiating the inhibitory effects of FASNi. Bars express mean ± SD. Statistical analyses were done using two-tailed unpaired Student’s t test, *p < 0.05, **p<0.01, ***p<0.001 and ****p<0.0001.

### FASN inhibition induces accumulation of PUFA-phospholipids in KMLC cells

To investigate the metabolic impact of FASN inhibition, we performed MS/MS^ALL^ untargeted lipidomic analysis. We found that FASNi induces accumulation of LysoPC only in KMLC cells. On the contrary, FASNi causes a concomitant decrease of TAG and an increase of DAG irrespectively of *KRAS* status (Fig. 4A and 4B). No significant changes were found in the activity of either the PC-specific phospholipase C (PLC) (Supplementary Fig. S5D) or the TAG-specific lipase ATGL (Supplementary Fig. S5E), suggesting FASNi does not affect PC and TAG lipolysis (Zechner et al., 2012). These findings suggest that KMLC uses *de novo* lipogenesis to fuel the synthesis of TAG and PC. Notably, we found that KMLC cells accumulate PUFA in LysoPC and PC upon FASNi (Fig. 4C-4F). In particular, LysoPC containing FA with two and four double bonds account for the LysoPC increase in FASNi-treated KMLC cells (Fig. 4C and 4D). Moreover, even though we did not detect a significant change in the total amount of PC (Fig. 4A and 4B), we observed the accumulation of PUFA-PC and depletion of SFA- and MUFA-PC in FASNi-treated KMLC cells ((Fig. 4E and 4F). This change in the composition of the acyl chains is specific for PC and LysoPC of KMLC, because other lipid classes, such as TAG, do not increase their incorporation of PUFA in response to FASNi (Fig. 4G and 4H). These data indicate not only that FASN provides SFA and MUFA for the synthesis of TAG and PC in KMLC, but also that its inhibition causes the incorporation of PUFA specifically in PC and LysoPC.

**Figure 4.**
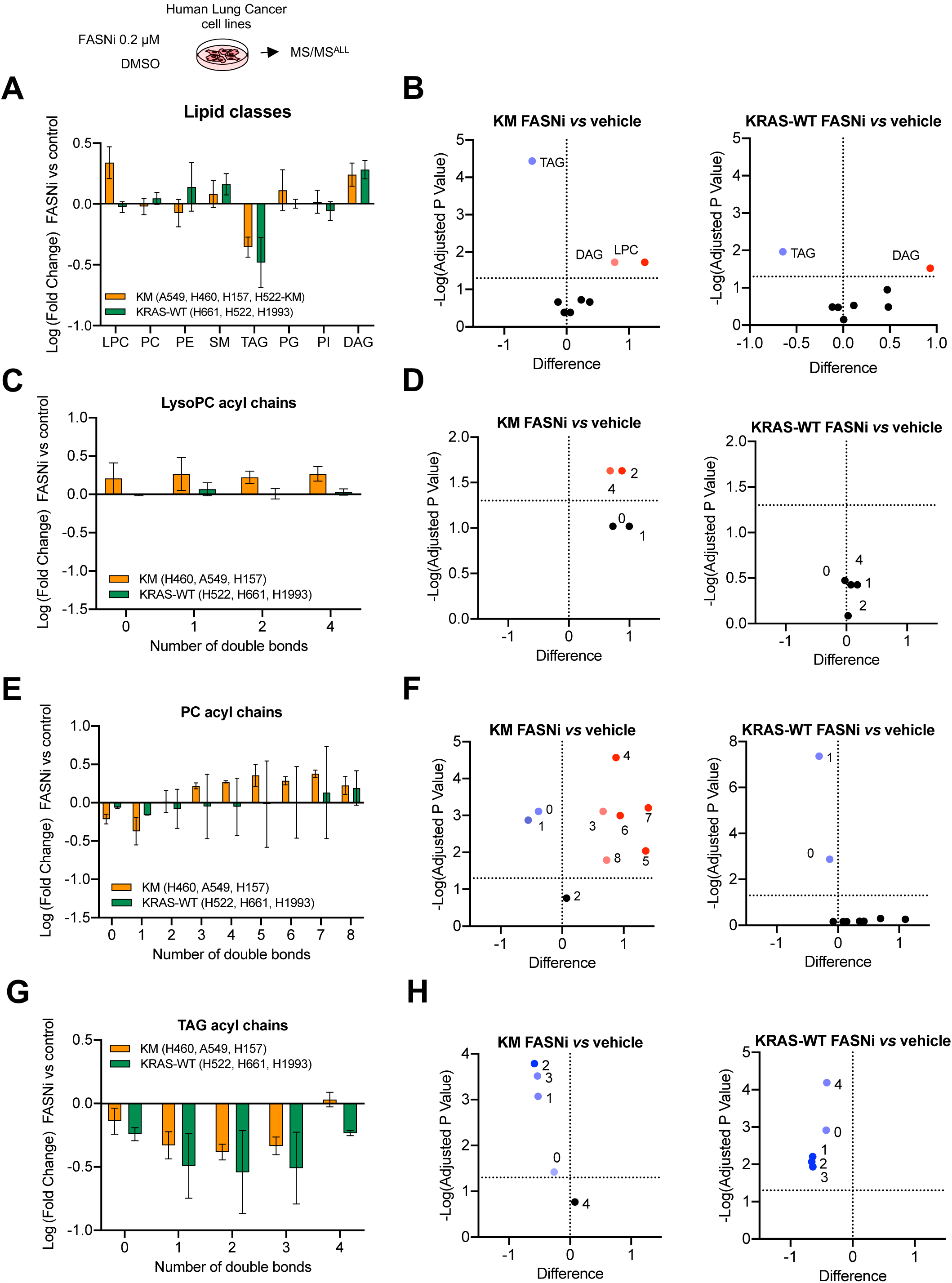
FASNi induces accumulation of PUFA-PC and PUFA-LysoPC in KMLC. **(A)** Variation of the lipid classes identified by MS/MS in KM and KRAS-WT LC cells. Bars represents Log (fold change) of FASNi treatment over vehicle control. **(B)** Volcano plots of multiple t tests representing the significant changes in lipid classes for the indicated comparisons (cutoff adj p<0.05). **(C-H)** Relative double bonds quantification in the indicated lipid classes and volcano plots showing the correspondent multiple t tests (cutoff adj p<0.05).

### The Lands cycle prevents the accumulation of PUFA in PC of KMLC

PC are synthetized via the Kennedy pathway which conjugates phosphocholine to DAG (KENNEDY and WEISS, 1956). However, the majority of PL synthesized via the Kennedy pathway are remodeled through the Lands cycle, which consists in the de-acylation of PC and re-acylation of LysoPC (Lands, 1960; Wang and Tontonoz, 2019; Zhao et al., 2008). To test whether FASN inhibition induces uptake and incorporation of exogenous PUFA, we used arachidonic acid (AA, FA 20:4) as PUFA proxy (Fig. 5A-D). In mammals, AA is provided by exogenous dietary sources rich either in AA or its parent molecule linoleic acid (LA, FA 18:2), which is desaturated and elongated in the endoplasmic reticulum to yield AA (Hanna and Hafez, 2018). Even though FASNi increases the total amount of AA in KMLC cells (Fig. 5A), ethyl acetate-2-^13^C metabolic flux analysis revealed a decreased isotope incorporation in AA of FASNi-treated KMLC cells (Fig. 5B). These data suggest that the intracellular pool of PUFA is dependent on their uptake from the microenvironment. This conclusion is consistent with our MALDI imaging data showing that PUFA-PC are enriched in the lung parenchyma surrounding the KM tumors (Fig. 1D-E).

**Figure 5.**
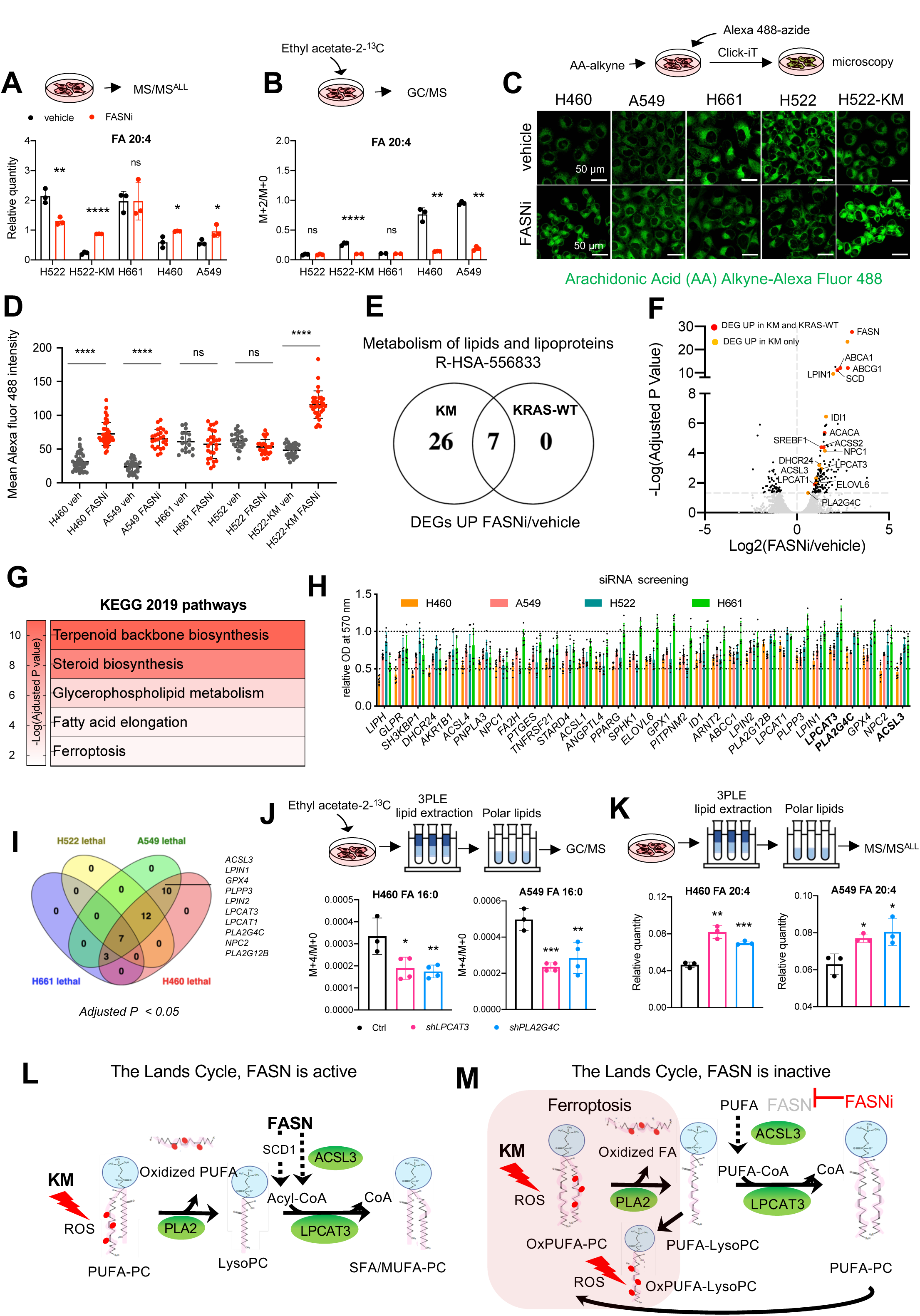
FASN and the Lands cycle limit PUFA content of phospholipids of KMLC. **(A, B)** Quantification of total (A) and newly synthetized (B) arachidonic acid (FA 20:4) in the PL fraction of the indicated cell lines. Tracer incorporation was measured after 7-hour incubation with Ethyl Acetate-1,2 ^13^C_2_. **(C, D)** Incorporation of arachidonic acid (AA) alkyne in the indicated cell lines treated as indicated, and its quantification. n=20-42 cells/sample. (**E)** Venn Diagram of the “metabolism of lipids and lipoproteins” (R-HSA-556833) genes upregulated in KM (H460, A549) and KRAS-WT (H661, H522) LC cells treated with FASNi. (**F, G**) Gene expression volcano plot and top KEGG pathways specifically upregulated in KMLC cells upon FASNi treatment. (**H, I**) Cell viability after siRNA-mediated knockdown of the indicated genes. Venn diagram summarizes lethal genes specific for KMLC cells (H460. A549, Dunnett’s multiple comparison test with cutoff adj P<0.05). **(J, K)** Quantification of newly synthetized-palmitate (FA 16:0) and total arachidonic acid (FA 20:4) in the PL fraction of indicated cell lines after shRNA-mediated knockdown of lysophosphatidylcholine acyltransferase 3 (*LPCAT3*) or phospholipase A2 group IVC (*PLA2G4C*). **(L, M)** Working model explaining the role of FASN in the regulation of the Lands cycle in KMLC. FASN is active: KM induces ROS that oxidize the PUFA acyl chain on PC. PLA2 removes the oxidized fatty acid (FA) on PC synthetizing a LysoPC. FASN and SCD1 produce saturated FA (SFA) and MUFA, respectively. SFA/MUFA are transferred to CoA by ACSL3. These acyl-CoAs are used by LPCAT3 to re-acylate the LysoPC forming again PC. Inhibition of FASN (J) causes the depletion of SFA/MUFA and uptake of exogenous PUFA for the re-acylation of LysoPC. This process increases the amount of PUFA-PC and PUFA-LysoPC which are oxidized under oxidative stress (oxPUFA-PC and oxPUFA-LysoPC). Accumulation of these lipid species leads to cell death via ferroptosis. Bars represent mean ± SD. In (A,B), (D), (J, K) Statistical analyses were done using two-tailed unpaired Student’s t test with ns, not significant, *p < 0.05, **p<0.01 and ***p<0.001.

To confirm this conclusion, we used click chemistry to conjugate AA-Alkyne to the Alexa Fluor 488-azide (Gaebler et al., 2013; Gao and Hannoush, 2014; Robichaud et al., 2016). We demonstrated that, even though the baseline AA uptake is lower in KMLC as compared to KRAS-WT cells, FASNi treatment increases the incorporation of AA only in KMLC cells (Fig. 5C and 5D). Noteworthy, ectopic expression of KM significantly increases AA uptake during FASNi treatment (H552-KM, Fig. 5C and 5D), indicating that this process is dependent on KM.

FASNi treatment upregulates several genes involved in the metabolism of lipids and lipoproteins in both KM and wtKRAS LC cells (Fig. 5E; Supplementary Table S3). However, we found that only FASNi-treated KMLC cells selectively upregulate metabolic genes such as *PLA2G4C*, *LPCAT3* and *ACSL3* (Fig. 5F). These genes are involved in the remodeling of phospholipids through the Lands cycle and in ferroptosis (Fig. 5G), a regulated form of cell death characterized by lipid peroxidation (Dixon et al., 2012; Lands, 1960). In particular, *PLA2G4C* is a member of the PLA2 family, which hydrolyses PL to produce free FA and lysophospholipids (LysoPL). LPCAT3 inserts acyl groups into LysoPL, specifically forming PC and PE (Zhao et al., 2008) and it is also required for cells to undergo ferroptosis (Dixon et al., 2015; Kagan et al., 2017). ACSL3 converts free long-chain FA into fatty acyl-CoA esters which undergo ß-oxidation or incorporation into PL (Yao and Ye, 2008). ACSL3 is required for KMLC tumorigenesis (Padanad et al., 2016), it plays an important role in AA metabolism in LC (Saliakoura et al., 2020) and protects cells from ferroptosis (Magtanong et al., 2019; Ubellacker et al., 2020). To validate these RNA-seq data, we perform a custom siRNA screen targeting 29 lipid metabolism genes. We found that 10 genes, including *PLA2G4C*, *LPCAT3*, *LPCAT1*, *ACSL3* and glutathione peroxidase 4 (*GPX4*), which counteracts ferroptosis by catalyzing the reduction of peroxided PL (Liu et al., 2018; Viswanathan et al., 2017; Yang et al., 2014, 2016; Zheng and Conrad, 2020), are selectively required for the viability of KM but not of KRAS-WT LC cells (Fig. 5H and 5I).

Consistently, KMLC cells are more sensitive than KRAS-WT LC cells to ML162, a specific inhibitor of GPX4 and an inducer of ferroptosis (Supplementary Fig. S6A and S6B), and we rescued the effect of FASNi using the anti-ferroptotic molecule ferrostatin-1 (Fer-1) (Supplementary Fig. S6A-S6F). Stable knock-down of *GPX4, LPCAT3* and *PLA2G4C* phenocopies the effect of ML162 or FASNi, inducing G2/M cell cycle arrest specifically in KMLC cells (Supplementary Fig. S6G and S6H), a feature often associated with ferroptosis (Greenshields et al., 2017; Lin et al., 2016). These results, along with the lipidomic profile of KMLC (Fig. 1; Fig. 4), indicate that the *de novo* lipogenesis is necessary to repair/prevent lipid peroxidation in KMLC by feeding the Lands cycle with SFA and MUFA. To demonstrate this hypothesis, we performed stable knock-down of *LPCAT3* and *PLA2G4C* in KMLC cells. Then we subjected them to 3PLE extraction (Vale et al., 2019) coupled with either ethyl acetate-2-^13^C metabolic flux analysis (Fig. 5J) or steady-state MS/MS^ALL^ lipidomic analysis (Fig. 5K). We determined that silencing either *LPCAT3* or *PLA2G4C* phenocopies FASN inhibition, decreasing the incorporation of *de novo* synthesized SFA, such as palmitate (FA 16:0) (Fig. 5J), while increasing the AA (FA 20:4) content of PL (Fig. 5K). These data demonstrate that both *de novo* FA synthesis and PL remodeling are necessary to prevent PUFA accumulation and ferroptosis in KMLC cells (Fig. 5L and 5M).

### FASN and the Lands cycle are required to deflect ferroptosis in KMLC

Using the lipid peroxidation probe C11-BODIPY (581/591), we confirmed that FASNi causes ferroptosis specifically in KMLC cells, leaving KRAS-WT cells unaffected (Fig. 6A and 6B). Notably, ectopic expression of *KM* in KRAS-WT cells causes C11-BODIPY oxidation in response to FASN inhibition (Fig. 6C and 6D), indicating that KM is required for the induction of ferroptosis. *TP53, STK11* and KEAP1/NFE2L2 (also known as NRF2) are frequently mutated in KMLC, deregulating cellular metabolism and oxidative stress (Galan-Cobo et al., 2019; Hassannia et al., 2019; Hayes and McMahon, 2009; Romero et al., 2017; Tarangelo et al., 2018). We did not find any correlation between their mutational status and sensitivity to FASNi in the LC cell lines we used (Supplementary Table S5). Therefore, these data indicate that the susceptibility of KMLC to ferroptosis depends on KM, rather than on *KEAP1*/*NFE2L2*, *TP53* and *STK11* co-occurring mutations.

**Figure 6.**
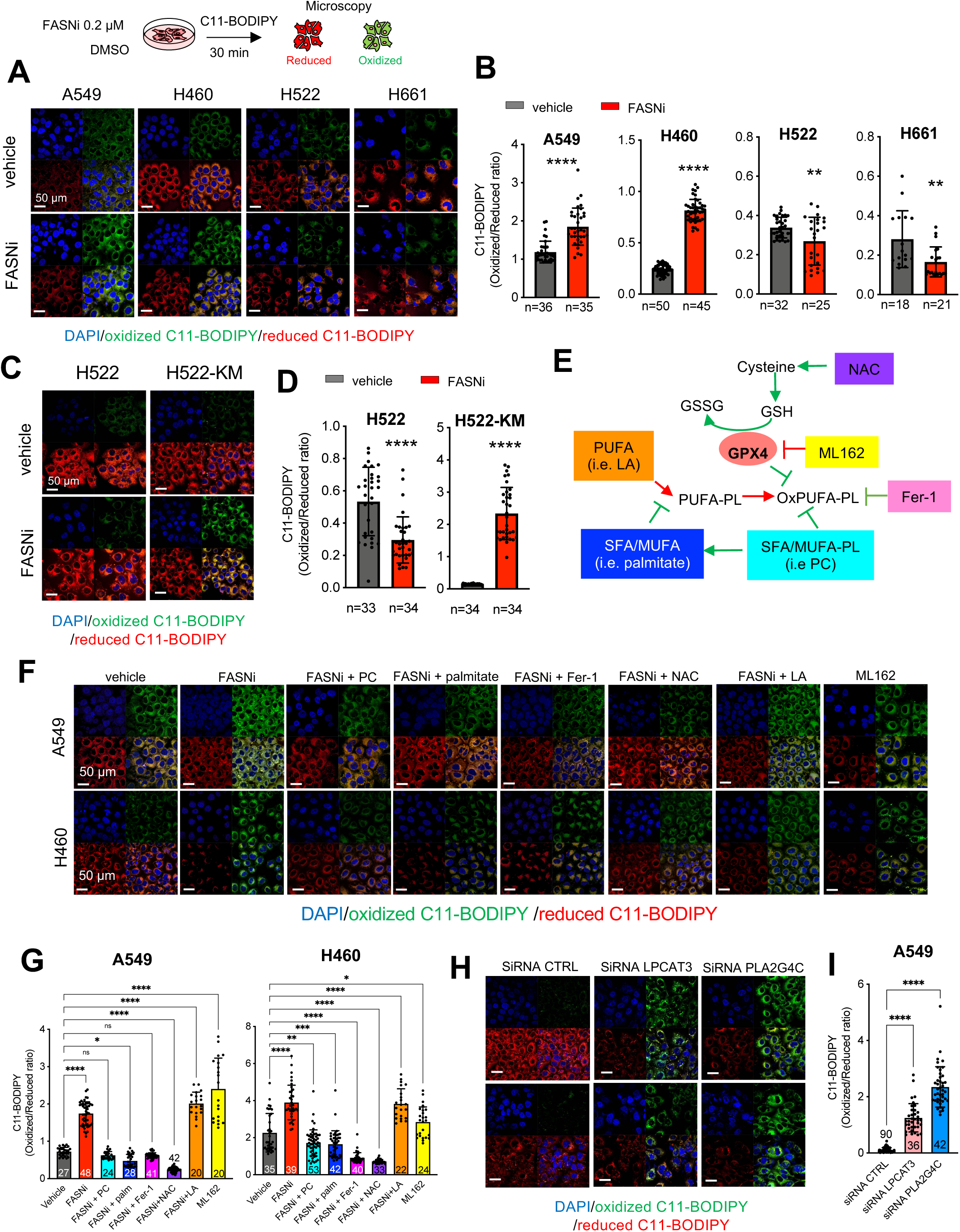
FASN and the Lands cycle are required to deflect ferroptosis in KMLC. **(A, D)** C11-BODIPY staining in KM, KRAS-WT, and H522-KM LC cells. Oxidized and reduced C11-BODIPY are indicated in red and green, respectively. Bars indicate the relative C11-BODIPY oxidation (n, number of cells). **(E)** Schematic of the GPX4 axis of ferroptosis and some of its regulators. Red=pro-ferroptosis; Green=anti-ferroptosis. NAC, N-acetyl cysteine; GSH, glutathione; GSSG, glutathione disulfide; ML162, GPX4 inhibitor; Fer-1, ferrostatin-1; PUFA, Polyunsaturated fatty acids; OxPUFA, oxidized PUFA; PL, phospholipids; MUFA, monounsaturated fatty acids; SFA, saturated fatty acids; LA, linoleic acid; PC, phosphatidylcholine. **(F, G)** C11-BODIPY rescue experiments in the indicated cell lines and their quantification. **(H, I)** C11-BODIPY stain on A549 cell line transfected with the indicated siRNAs (48hrs post transfection) and their quantification. Numbers in the bars indicate number of cells. Bars represent mean ± SD. In (B) and (D) two-tailed unpaired student t test, in (G) multiple t tests with ns, p>0.05; *p<0.05; **p<0.01; ***p<0.001; ****p<0.0001.

Next, we used the 3PLE extraction method coupled with UV absorbance to measure the FA oxidation in the neutral (*e.g.* TAG) and polar lipid (*e.g.* PL and LysoPL) fractions of KMLC cells (Supplementary Figure S7A) (Vale et al., 2019). Indeed, as oxidation occurs in lipids containing two or more double bonds, it causes an increase in the UV emission that can be used to quantify the rate of primary lipid oxidation (Kim and LaBella, 1987). We found that FASNi induces oxidation specifically of the polar lipid fraction of KMLC cells (Supplementary Figure S7B), without perturbing neutral lipids (Supplementary Figure S7C). This finding is consistent with the observation that in KMLC cells, FASNi leads to an enrichment of PUFA specifically in PC and LysoPC, but not in TAG (Fig. 4). Accordingly, PC supplementation quenches ROS production and propagation in KMLC cell treated with FASNi (Supplementary Figure S7D and Supplementary Video).

To further demonstrate that FASN and the Lands cycle are necessary to prevent ferroptosis, we treated KMLC cells with FASNi and/or molecules targeting the cysteine/GSH/GPX4 system, one of the mainstays restricting ferroptosis (Fig. 6E). Noteworthy, while PC, palmitate, ferrostatin-1 (Fer-1), and N-acetyl-cysteine (NAC) rescue the C11-BODIPY oxidation induced by FASNi, the PUFA linoleic acid (LA) does not (Fig. 6F, G). Furthermore, silencing of *LPCAT3* or *PLA2G4C* induces significant C11-BODIPY oxidation in KMLC cells (Fig. 6H, I).

### Pharmacologic inhibition of FASN suppresses KMLC *in vivo*

We explored the preclinical significance of our findings using transgenic *TetO-Kras^G12D^* mice (Fig. 7A, B) and xenografts of A549 and H460 KMLC cells (Fig. 7C, D). In all the preclinical mouse models, FASNi causes a potent anti-tumor effect without overt systemic toxicities (Fig. 7A-D; Supplementary Fig. S8A-C), and it depletes intratumor and serum palmitate (Supplementary Fig. S8D). Notably, FASNi induces lipid oxidation in both autochthonous and xenograft KMLC, as demonstrated by C11-BODIPY staining (Fig.7F-7H). In addition, lipidomic analysis (Fig. 7H) and MALDI imaging (Supplementary Fig. S8E) confirmed that inhibiting *de novo* lipogenesis causes accumulation of PUFA specifically in PC and Lyso-PC of KMLC tumors. On the other hand, TAG are downregulated and DAG are upregulated, independently of the saturation of their acyl chains (Fig. 7H; Supplementary Fig. S8E). These data phenocopy the findings obtained *in vitro* (Fig. 4) providing further evidence that FASNi induces ferroptosis in KMLC. This preclinical evidence will inform the ongoing clinical trial with the human specific FASN inhibitor TVB-2640, which is showing promising results in KMLC patients (NCT03808558, Fig. 7I).

**Figure 7.**
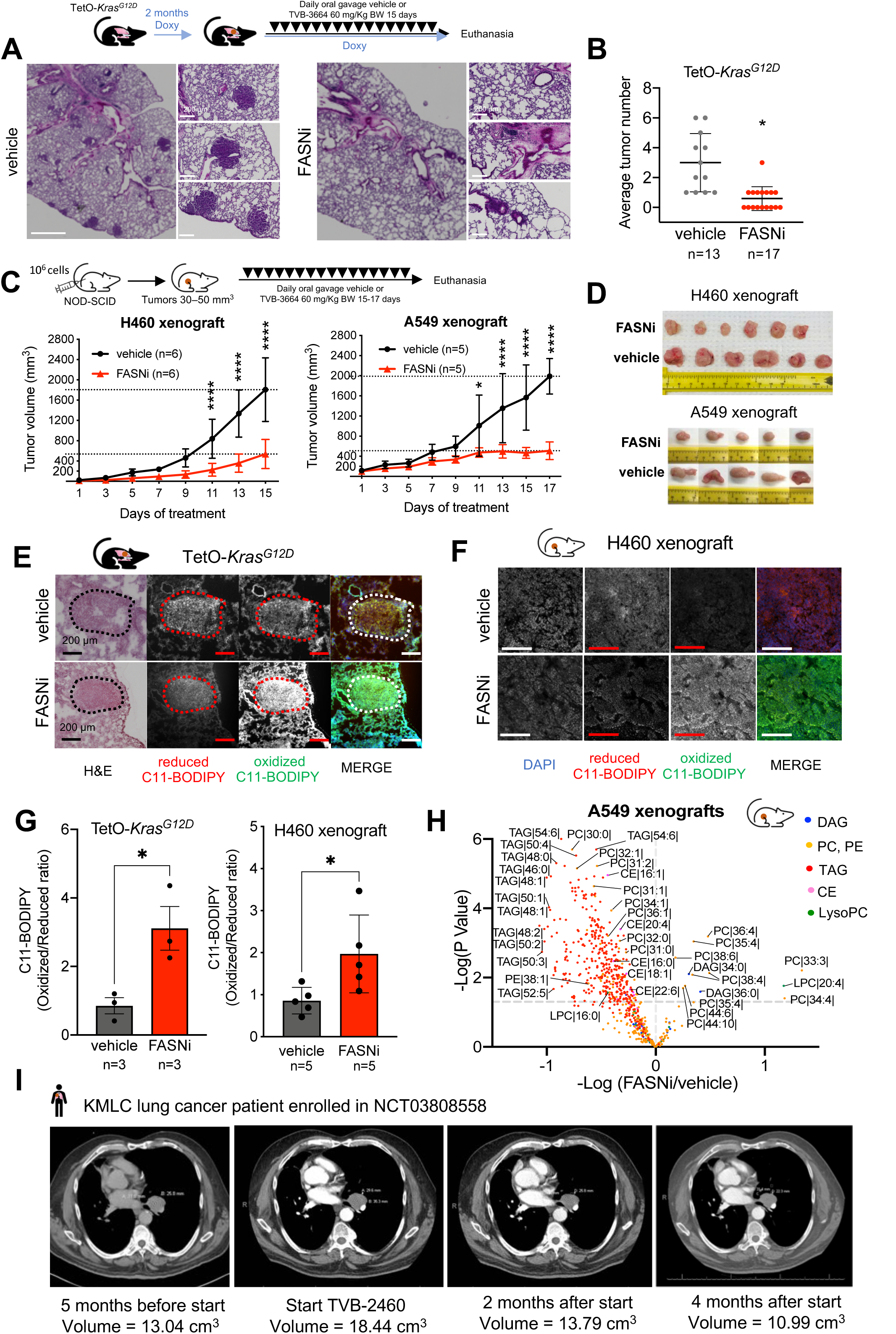
FASNi is effective in KMLC *in vivo.* **(A, B)** Representative H&E pictures of the lungs of TetO-*Kras* mice treated as indicated and tumor burden quantification. n=number of mice. **(C, D)** *In vivo* growth curves and post-resection pictures of H460 and A549 xenografts in NOD/SCID mice treated as indicated. **(E-G)** Representative pictures of C11-BODIPY staining of the lungs of TetO-*Kras* and of H460 xenografts, and their quantification. Dotted circles in (E) indicate lung tumor. **(H)** Volcano plot showing lipid species identified by MS/MS differentially represented in FASNi *versus* vehicle (n=5 mice/group). P values and difference were calculated using multiple t tests (p<0.05). **(I)** Serial CT-scans of a representative lung cancer patient enrolled in the NCT03808558 clinical trial with the FASNi derivative TVB-2640 (modified to improve pharmacokinetic properties in humans). Note the 30% reduction of estimated tumor volume after 8 weeks of treatment.

## Discussion

KM is the most commonly mutated oncogene in cancer (Prior et al., 2020). There is a considerable interest in determining the mechanisms underlying KM-driven tumorigenesis and KM-dependent tumor survival. This knowledge is a prerequisite to develop novel therapies as well as to optimize the use of existing drugs that target either KM or its downstream signaling pathways. It is well know that KM contributes to the regulation of cancer metabolism; however, no metabolic network has been established as *a bona fide* therapeutic target (Boroughs and DeBerardinis, 2015). Furthermore, it is also emerging that metabolic adaptations mediate resistance to KRAS inhibitors (Santana-Codina et al., 2020).

Here, we report that KMLC depends on *de novo* FA synthesis and PL remodeling to cope with oxidative stress and to escape ferroptosis. In KMLC, KM upregulates FASN, whose inhibition causes lipid peroxidation and ferroptosis. These effects are specific since metabolites immediately downstream FASN (i.e. palmitate or PC) rescue this phenotype. We determined that KM, but not EGFR-MUT, is necessary and sufficient to establish the dependency on FASN. On the contrary, we found no correlation with *TP53*, *STK11/LKB1* or *NFE2L2/NRF*2 which are frequently co-mutated with KM and known to influence metabolism and oxidative stress (Galan-Cobo et al., 2019; Romero et al., 2017).

It is a long-standing observation that *FASN* is overexpressed in several cancer types and that KM upregulates its expression (Gouw et al., 2017). It has been proposed that FA meet the requirements of highly proliferative cells providing building blocks for membranes, or sustaining ATP production through ß-oxidation (Baenke et al., 2013). However, the inhibition of these processes does not affect KRAS-WT cells and does not explain the exquisite and specific dependency of KMLC on *de novo* lipogenesis.

Our comprehensive metabolic and functional analysis shows, for the first time, that KM upregulates FASN to increase the synthesis of SFA and MUFA to prevent and repair lipid peroxidation in LC. Our conclusion is supported by unprecedented metabolic flux analysis showing how FASNi induces the incorporation of PUFA into PL, in particular PC. When *de novo* lipogenesis is impaired, KMLC increases the uptake of exogenous PUFA. However, even though we demonstrated that cellular lipolysis does not account for this process, we cannot exclude that other mechanisms like lipophagy and macropinocytosis (Kamphorst et al., 2013b; Kounakis et al., 2019) might contribute to the enrichment of PUFA observed after FASN inhibition.

Because of their multiple double bonds, PUFA are the major substrate of lipid peroxidation (Choi et al., 2011; Dixon et al., 2012; Hassannia et al., 2019). Therefore, cells use the Lands cycle to limit the amount of PUFA incorporated in PL, reducing lipid peroxidation and the likelihood to trigger ferroptosis (Pérez et al., 2006; Saliakoura et al., 2020; Surette et al., 1999). Accordingly, we found that silencing key regulators of the Lands cycle induces PUFA accumulation in the side chains of PL, leading KMLC to ferroptosis.

KM expression and/or activation of the RAS/MAPK pathway have been reported to sensitize cells to ferroptosis inducers (Poursaitidis et al., 2017; Yagoda et al., 2007; Yang and Stockwell, 2008). However, later studies challenged these conclusions (Yang et al., 2014). We reason that this incongruency is explained by the fact that activation of KRAS pathway increases the susceptibility to ferroptosis, but that cell context specific factors contribute to its execution.

Our data indicate that KMLC has a lipidomic profile that is reminiscent of alveolar type II cells (AT2), which are considered the cells of origin of LC (Rowbotham and Kim, 2014). AT2 cells are a major source for surfactant lipids secreted into the alveoli. Surfactant lipids comprise about 90% of pulmonary surfactant, a lipoprotein complex which decreases surface tension in the post-natal lung and prevents alveolar collapse. PC are the most abundant surfactants lipids (Griese et al., 2015). Notably, SFA and MUFA-PC like dipalmitoylphosphatidylcholine PC 32:0 (DPPC, 16:0/16:0), PC 32:1 (16:0/16:1) and PC 30:0 (16:0/14:0) are the most abundant members of surfactant PC both in AT2 cells and in KMLC (Batenburg, 1992; Holm et al., 1996; Kyle et al., 2018). Thus, it seems reasonable to conclude that KM hijacks lung specific mechanisms evolved to mitigate oxidative stress and ferroptosis under the high oxygen tension conditions found in the pulmonary alveolus (Guo et al., 2019). Indeed, transitioning from the fetal hypoxic environment to air breathing at birth requires the activation of adaptive responses, as demonstrated by the observation that, mouse AT2 not only have high expression of *Kras* soon after birth, but they also exhibit high expression of genes involved in the unfolded protein response (UPR), antioxidant response and lipid synthesis (Guo et al., 2019). Hence, FASNi interferes with the PC-dependent antioxidant function, decreasing the threshold to ferroptosis in the context of high ROS in KMLC. In this scenario, we reason that the PUFA-rich lung environment, surrounding KMLC *in vivo*, may represent a “ticking bomb” for KM tumors. This is in agreement with our observation that exogenous palmitate bypasses FASNi-induced cell death and with previous studies showing that exogenous MUFA confer resistance to ferroptosis (Magtanong et al., 2019). Future studies are needed to determine whether FA deriving from dietary sources or the tumor microenvironment could directly influence KMLC tumorigenesis and response to therapy.

Our data predict not only that FASNi will be effective in KMLC therapy, but also that other inducers of ferroptosis may exert selective anti-tumor effects in KMLC. It is noteworthy that TVB-2640, a FASNi derivative with improved pharmacokinetic properties, is showing promising results in a phase II trial in KMLC (NCT03808558) and that several other FDA-approved drugs (*e.g.* sorafenib, sulfasalazine, artesunate, lanperisone) are known to induce ferroptosis in certain cancers (Eling et al., 2015; Gout et al., 2001; Li et al., 2020; Louandre et al., 2013, 2015; Shaw et al., 2011; Zhang et al., 2019). Therefore, it is likely that cancers other than KMLC rely on FASN and the Lands cycle to overcome ferroptosis and that this dependency could be exploited in the clinic.

## Supporting information

Supplementary Table S1_List of Human LC samples and PDXs

Supplementary Table S2_siRNA screening

Supplementary Table S3_YTMA310 and TMA4

Supplementary Table S4_RNA-seq DEGs KM and KRAS-WT LC cell lines

Supplementary Table S5_Mutations NSCLC cell lines

## Acknowledgements

The preliminary part of this work was done at UT Southwestern medical Center. CPRIT RP140672, CPRIT RP1606552, The University of Cincinnati College of Medicine (PPS). Lung Cancer SPORE (P50CA70907) (JDM, PPS, JWS, KH, IIW), the Harold C. Simmons Cancer Center through NCI Cancer Center support grant and 2P30CA016672. Cancer Prevention and Research Institute of Texas CPRIT RP160652 The University of Texas MD Anderson Cancer Center. YTMA 310 was funded in part by the Yale SPORE in Lung Cancer P50 CA 196530 (PI: Roy Herbst). We thank Dr. Monte Winslow for kindly providing the murine LSL-*KRAS^G12D^* cell lines, John M. Shelton for helping set up LMD conditions, Dr. Ken Greis and Dr. Robert Ross at UC Proteomics and Metabolomics Laboratory to provide guidance and instrumentation for the HPLC-MS/MS experiments, and Dr. Peter Pathrose for performing KRAS mutation analysis on human lung cancer samples.

## Author contribution

CB and CA equally contributed to this study; CB, CA and PPS designed the study; CB, CA, GVDV, MM, ACC, DB performed experiments; SB and CA performed bioinformatic analysis; GVDV and JM gave technical assistance with lipidomic experiments; JM, JDM, KP, BG provided some key reagents and resources; GK provided TVB-3664 and TVB-2632; SLS provided the human lung cancer sample; DG provided CT-scans from NCT03808558 trial; KH provided pathology assistance with YTMA 310; DL performed immunohistochemistry analysis on TMA 4; MGR, and LSS provided tissue acquisition and pathology assistance with TMA 4; CBe clinical database management and HK mutational database acquisition and management (TMA 4); CA, CB and PPS wrote the manuscript with comments from all authors.

## Data availability

RNA-seq data were deposited in GEO under the accession number GSE168782.

All data supporting the findings of this work are available within the article and Supplementary Information. Further requests for resources, reagents and data should be directed to and will be fulfilled by the corresponding author.

## Conflict of interest statement

The authors declare no potential conflicts of interest.

## Notes

### Competing Interest Statement

The authors have declared no competing interest.

