## Supplementary figures and images for "Targeting *de novo* lipogenesis and the Lands cycle induces ferroptosis in KRAS-mutant lung cancer"

### Supplementary Video 1

#### Slide 1
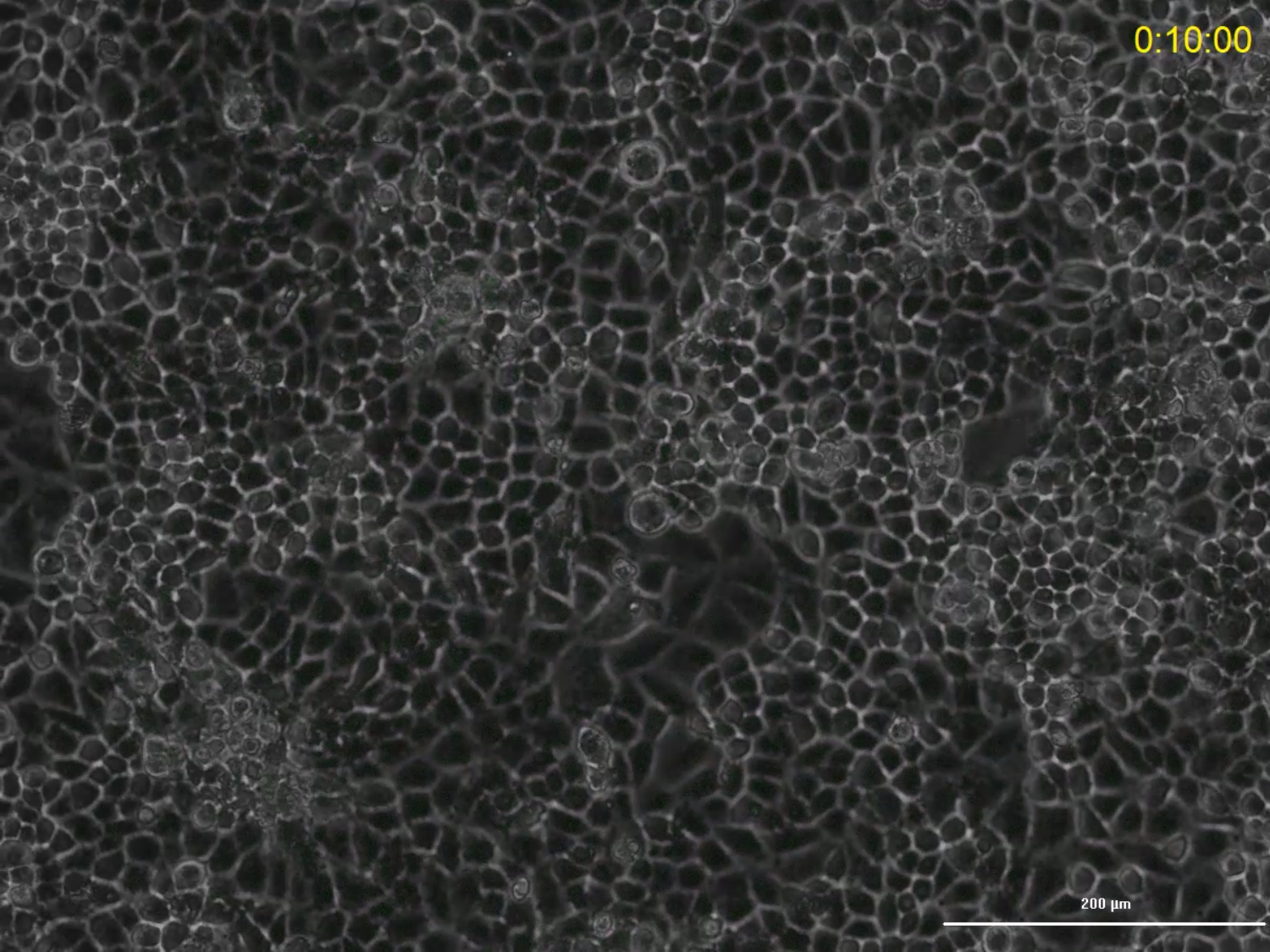

### Supplementary video 2

#### Slide 1
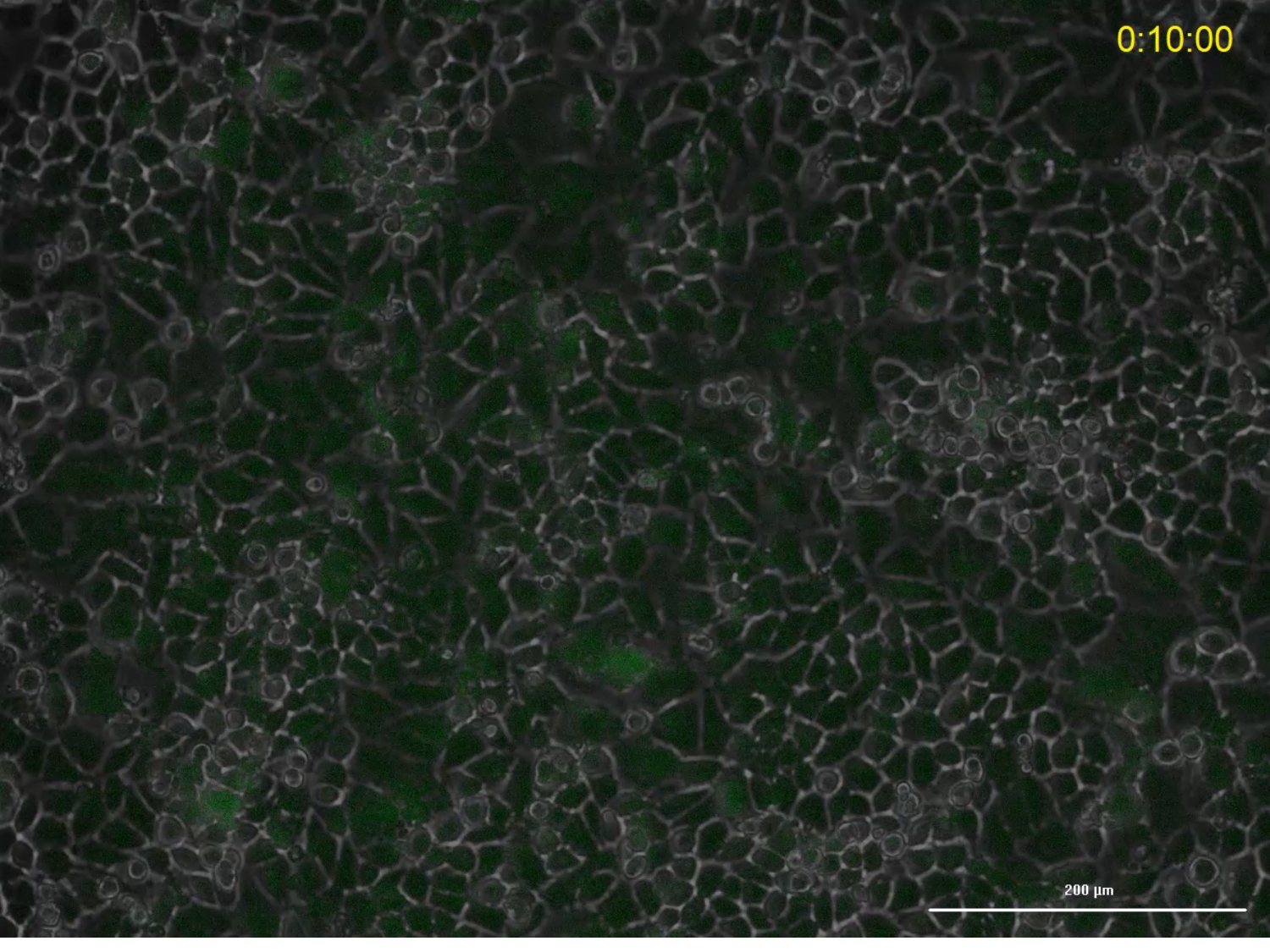

### Supplementary video 3

#### Slide 1
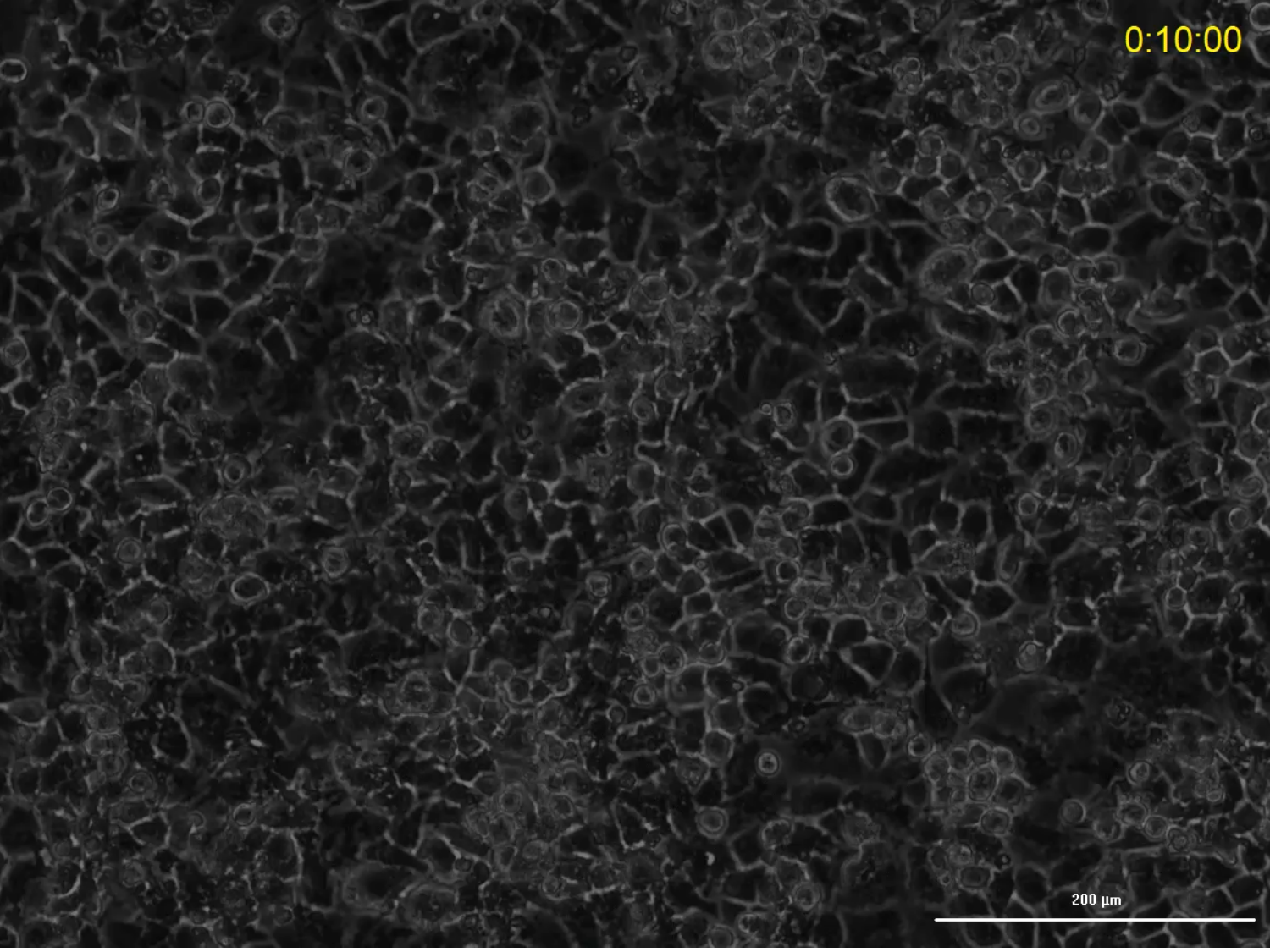
