## Supplementary information for "Targeting *de novo* lipogenesis and the Lands cycle induces ferroptosis in KRAS-mutant lung cancer"

**Affiliations:**

**Running title:** Mutant KRAS lung cancer depends on FASN and the Lands cycle

**Conflict of interest statement:** The authors declare no potential conflicts of interest.

**Content**

Supplementary methods

Supplementary Figures S1-S8

**Other Supplementary Materials for this manuscript include the following:**

Supplementary Tables S1 to S5 as Excel files

Supplementary Videos S1-S3

**Supplementary methods**

*Detection of FASN in human KMLC*. We obtained data regarding FASN expression in human KM, EGFR-MUT, KRAS-WT and EGFR-WT LC cell lines from the Cancer Cell Line Encyclopedia (CCLE) database (Barretina et al., 2012), www.broadinstitute.org/ccle).

YTMA-310 tumor tissue microarray (TMA), Yale Pathology Tissue Service (YPTS), consists of 30 EGFR-mutant, 43 KRAS-mutant, 66 EGFR and KRAS wild-type (EGFR/KRAS WT) NSCLC patient samples, and 6 cell line cores (HCC193, A431, HT29, SW480, MCF7, and H2126) (Toki et al., 2017, 2018). TMA-4 was from Department of Translational Molecular Pathology at the University of Texas MD Anderson Cancer Center (n=242 adenocarcinoma specimens, Supplementary Table S3).

Sections were deparaffinized and subjected to antigen retrieval in Citrate buffer. After 1h antigen block in 2.5% normal horse serum (ImmPRESS$^{TM}$ reagent kit, Vector Laboratories), followed by overnight incubation with mouse FASN monoclonal antibody (sc-55580, SCBT, dilution 1:500 in 2.5% normal horse serum) at 4°C. Then slides were stained with HRP-conjugated anti-Mouse Ig (cat. MP-7402, ImmPRESS$^{TM}$ reagent kit, Vector Laboratories) and the peroxidase substrate kit, DAB (SK-4100, Vector Laboratories). Scoring was carried out blindly. Immunoreactivity score (0-300) was determined multiplying the staining intensity (0, no staining; 1, weak; 2, moderate; 3, strong staining) and the percentage of positive tumor cells in each core (0-100%).

*Preparation of BSA-Conjugated Palmitate*. 1mM Sodium Palmitate/0.17 mM BSA solution (6:1 molar ratio Palmitate:BSA) was prepared using ultra fatty acid free BSA (SIGMA ALDRICH, A6003-25G). Briefly, BSA was dissolved in 150 mM NaCl solution. Half of the solution was conjugated with Sodium Palmitate (SIGMA ALDRICH, P-9767), stirring 1 hour at 37°C; the remaining was used as vehicle control in subsequent assays.

*Drug Stability Assay*. We seeded 2000 H460 cells/well in 50 µL complete medium in a 96-well plate at 37°C overnight. The next day fresh RPMI + 5% FBS dialyzed medium with 200 nM TVB-3664 and added to the wells. Medium was harvested at different time points (t=0, 60 min, 240 min, 480 min, 24 hrs). Briefly, 100 µL of media was removed from each well and transfer to a tube containing 125 µL of organic crash solution (Methanol (MeOH), 0.2% Formic Acid (FA), 50 ng/mL internal standard). Wells were washed with 100 µL ice-cold PBS and PBS was transferred to the corresponding tube containing the crush solution. Cells were trypsinized, trypsin was blocked with 100 µL of cold medium and transferred to a new tube. Wells were rinsed with 125 µL of crush solution and the rinsed solution was transferred to the tube. Tubes were vortexed for 15 seconds and incubated at room temperature for 10 min, spun at 13200 rpm for 5 minutes. 190 µL of supernatant was transferred to an LCMS vial with insert and analyzed by LCMS (QTrap 4000). Ion Source/Gas Parameters: CUR = 310, CAD = Med, IS = 4500, TEM = 650, GS1 = 70, GS2 =70. Buffer A: Water + 0.1% FA; Buffer B: MeOH + 0.1% FA; flow rate 1.5 mL/min; column Agilent C18 XDB column, 5 micron packing 50 X 4.6 mm size; 0 - 1.0 min 5% B, 1.0 - 1.5 min gradient to 100% B, 1.5 - 2.5 min 100% B, 2.5 - 2.6 min gradient to 5% B, 2.6 - 3.5 5% B; IS: tolbutamide (transition 271.2 to 91.2); TVB-3664 transition 469.19 to 437.1.

*Measurement of intracellular drug concentration.* We plated H460 cells on 10 cm dishes at $5x10^5$/plate and incubated at 37°C overnight. The next day, media was removed and replaced with fresh RPMI + 5% FBS dialyzed medium containing TVB-3664 (200nM). Assay time points were 2, 4, 8 and 24 hrs. At each time point, media was removed, cells washed twice with ice-cold PBS and trypsinized. Cells were resuspended in PBS at $1x10^6$ cells/mL. After adding 250 µL of crash solution (MeOH, 0.2% Formic Acid, 0.05 ng/mL internal standard). After incubation at RT for 10min and centrifugation, the supernatant (160 µL) was analyzed by LCMS (QTrap 4000). QTrap

4000 parameters were the same as set for drug stability assay. To analyze data, blank signal was subtracted from all points on standard curve and then plotted in GraphPad to create a standard curve. The standard curve was used to quantitate samples.

*Phosphatidylcholine specific phospholipase C (PC-LPC )enzymatic activity.* PC-PLC activity in cellular and tumor lysates was assessed by EnzChek™ direct phospholipase C assay kit (Thermo Scientific, E10215) previously described (Upham et al., 2008). We used 50 μg of cell lysate, in triplicate.

*Lipase enzymatic activity*. We used a Lipase activity colorimetric assay (Abcam, ab102524) that quantifies glycerol deriving from TAG hydrolysis mediated by ATGL. Briefly, 2 x 106 cells corresponding to each experimental condition tested were analyzed in triplicate. Lipase was used as positive control, while samples without the reaction mix were used background controls.

*Determination of phospholipid oxidation by UV/VIS spectroscopy.* For estimation of conjugated dienes, lipids from cell samples were fractioned in the polar (phospholipids) and neutral fractions, using the 3PLE method (Vale et al., 2019). Lipid fractions were dried and dissolved in ethanol. For both fractions, the spectra of ultraviolet absorption between 200 nm and 300 nm were measured as previously described (Kim and LaBella, 1987).

*CellROX™ Green assay and live imaging*. Sub-confluent H460 cells in 6-well plates were treated with either vehicle, TVB-3664 (0.2 μM) or combination of TVB-3664 (0.2 μM) and 16:0-18:1 PC (50 μM). After 72 hrs, CellROX Green reagent was spiked in at final concentration of 5 μM, and incubated for 30 min at 37°C. Plates were imaged over 1 h with one image taken every 5 minutes. Time-laps imaging was conducted using Lionheart FX automated microscope (BioTek) and Gen5 software.

**Supplementary Figure legends**

**Supplementary Figure S1. Mutant EGFR LC and KMLC have distinct lipidomic profiles. (A)** H&E staining and MALDI imaging pictures of TetO-*EGFR*$^{L858R}$ lungs. Rainbow scale represents ion intensity normalized against the total ion count (%) and black circles indicate the major tumor areas.**(B)** HPLC-MS/MS analysis of EGFR-MUT (n=2) and KM (n=5) lung cancer patient-derived xenografts (PDXs). P values and difference (EGFR-MUT *versus* KM) were calculated using multiple t tests ($p<0.05$). **(C)** Proposed mechanisms of KM-dependent regulation of lipid synthesis in lung cancer. KM activates *FASN* and *ACSL3* which synthesize saturated fatty acids (SFA) and SFA-CoAs respectively. SFA-CoAs are either desaturated by SCD1 to monounsaturated FA (MUFA-CoAs) or directly used for the synthesis of TAG and PC.

**Supplementary Figure S2. FASN is a selective vulnerability of KMLC.** (**A, B**) Crystal violet and MTT viability assays of the indicated murine LSL-*Kras*$^{G12D}$ cell lines treated as indicated. $IC_{50}$, inhibitory concentration 50. (**C**) Cell cycle analysis on the indicated human LC cell lines treated as indicated. RPMI 5% FBS = regular medium; RPMI 5% dFBS= medium with dialyzed FBS; RPMI 0.02% DMSO = vehicle. (**D**) Crystal violet assay of the indicated cells transfected with the indicated siRNAs and correspondent immunoblot of FASN. NT, non-targeting. (**E**) Cell cycle analysis of KM and KRAS-WT LC cells transfected with the indicated siRNAs. siRNA FASN pool = siFASN #1-4. **(F)** Cell viability assay in EGFR-MUT LC cell lines treated as indicated for 7 days.

**Supplementary Figure S3. FASN is upregulated in KMLC. (A)** Immunoblot analysis of FASN in micro-dissected lung tumors of CC10rtTA; TetO-*Kras*$^{G12D}$ mice +/-doxy for the indicated number of days. **(B)** FASN expression in human non-small cell lung cancer (NSCLC) cell lines from the cancer cell line encyclopedia (CCLE) of the indicated genotype. TPM, transcript per million. n=number of cell lines. **(C, D)** Representative images of FASN immunostaining in LC specimens of

YTMA 310 and its H-score quantification. n=number of cores. **(E, F)** Representative images of FASN immunostaining in LC specimens of TMA 4 and its H-score quantification. n=number of cores. **(G)** *FASN* expression in human cell lung cancer cell lines from the cancer cell line encyclopedia (CCLE) of the indicated genotype. Data are expressed as mean ± SD. **(H)** FASN H-score quantification of EGFR-MUT and EGFR-WT cores in YTMA 310 (n=number of cores). In (B), (D), (F) and (G, H) two-tailed unpaired Student's t test with ns, not significant, *p < 0.05, **p<0.01, ***p<0.001 and ****p<0.0001.

**Supplementary Figure S4. KM expression is sufficient to induce dependency on FASN. (A)** Oil red O staining for lipid droplets in the H661 (KRAS-WT) parental cell line and H661 transduced with KM . **(B)** Crystal violet assay of the indicated cell lines treated with vehicle or FASNi (0.2 µM). **(C)** Crystal violet assay of H522-KM cells transfected with the indicated siRNAs and correspondent immunoblot of FASN. NT, not targeting.

**Supplementary Figure S5**. **FASNi uptake, stability and activity in KM and KRAS-WT LC. (A, B)** Intracellular accumulation and stability of FASNi in H522 (KRAS-WT) and H460 (KM) cell lines (n=3 replicates) assessed by LC/MS. $t_{1/2}$= drug half-life. **(C)** Quantification of newly synthesized FA via GC/MS in A549 cells treated as indicated. M+2/M+0 ratio is reported. Palmitate, FA 16:0; palmitoleate, FA 16:1n7; stearate, FA 18:0; vaccenate, FA 18:1n7; oleate, FA 18:1n9 (n=3 independent experiments). **(D)** Oil red O staining for detection of lipid droplets (arrows) in the indicated cell lines. **(E, F)** Relative activity of PC-specific phospholipase C (PC-PLC) and of adipose triglyceride lipase (ATGL) in the indicated LC cell lines. Data are expressed as mean ± SD (A, B-D, G, H). Statistical significance was assessed using Student's t test with ns, p>0.05, not significant, **p<0.01, ***p<0.001 and ****p<0.0001.

**Supplementary Figure S6. KM expression lowers the threshold to ferroptosis induction. (A, B)** MTT viability assay of the indicated KM and KRAS-WT LC cells treated with increasing concentrations of ML162 (GPX4 inhibitor). $IC_{50}$ values are represented in the bar graph. **(C, D)** MTT viability assay of A549 and H522-KM cells treated with FASNi or ML162 in combination with increasing concentration of the ferroptosis inhibitor Ferrostatin-1. **(E, F)** Cell cycle analysis of the indicated cell lines treated as reported or transfected with the indicated inducible shRNAs. OFF/ON = -/+ doxy. **(G)** RNA expression of *GPX4, LPCAT3* and *PLA2G4C* before or after induction of the shRNAs. Bars represent mean ± SD. In (C, D) unpaired two-tailed student t test with ns, not significant, *$p < 0.05$, **$p<0.01$ and ***$p<0.001$.

**Supplementary Figure S7. FASNi causes phospholipid peroxidation. (A-C)** Schematic workflow and quantification of lipid peroxidation by UV absorbance in the polar and neutral lipid fractions of the indicated KMLC cell lines. **(D)** Representative time lapse snapshots of ROS production in H460 treated as indicated. CellROX green accumulation indicates oxidative stress. See also Supplementary videos 1-3.

**Supplementary Figure S8. Body weight, palmitate quantification and MALDI imaging of KMLC mouse models. (A-C)** Body weight of TetO-*Kras* and xenograft-bearing mice treated as indicated. **(D)** Palmitate quantification in xenografts and sera of NOD/SCID mice (n=5 mice/group). **(E)** Representative MALDI images of the indicated lipid species in A549 xenografts. Rainbow scale represents ion intensity normalized against the total ion count (%).

### Supplementary Figure 1

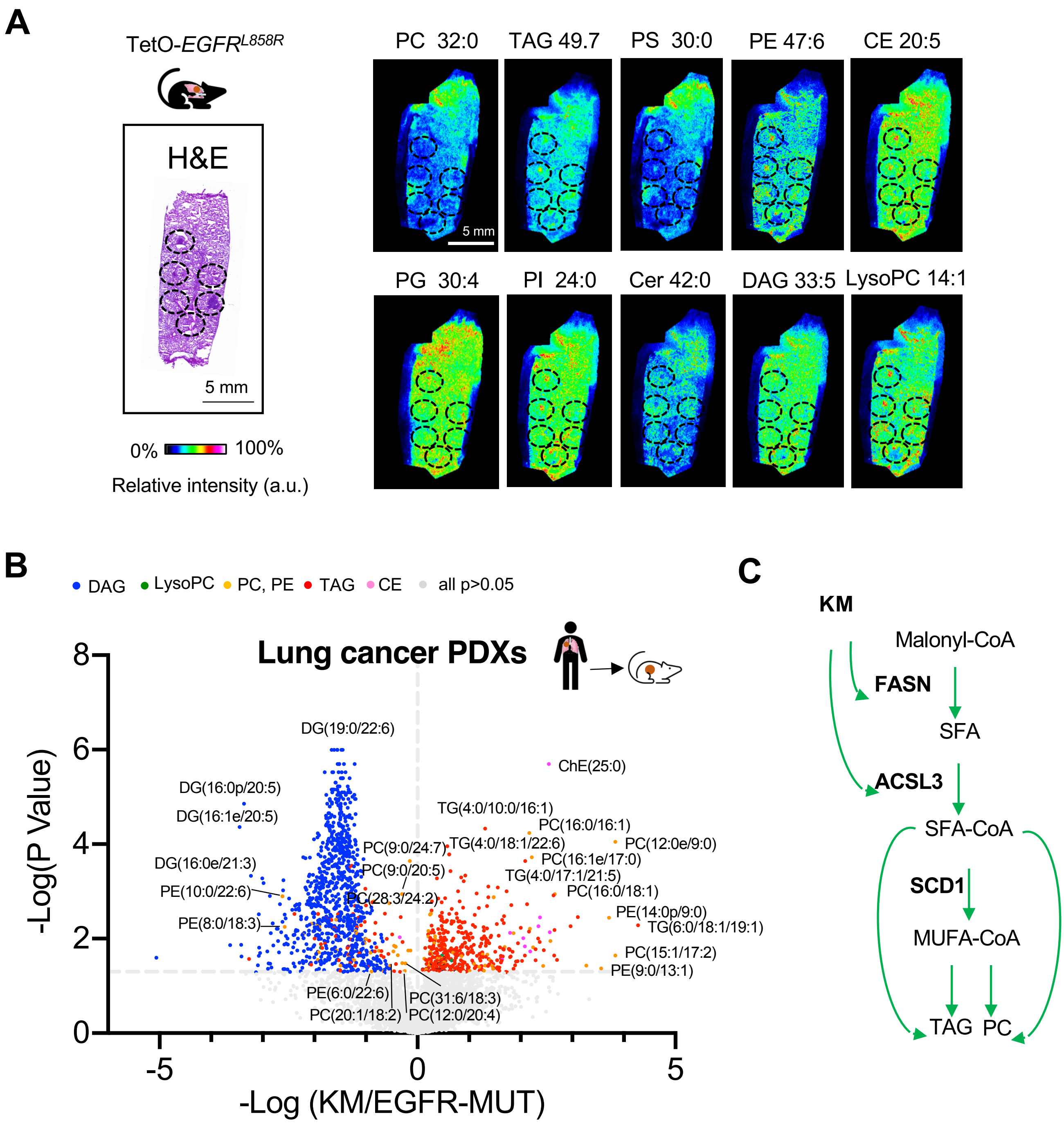

### Supplementary Figure 2

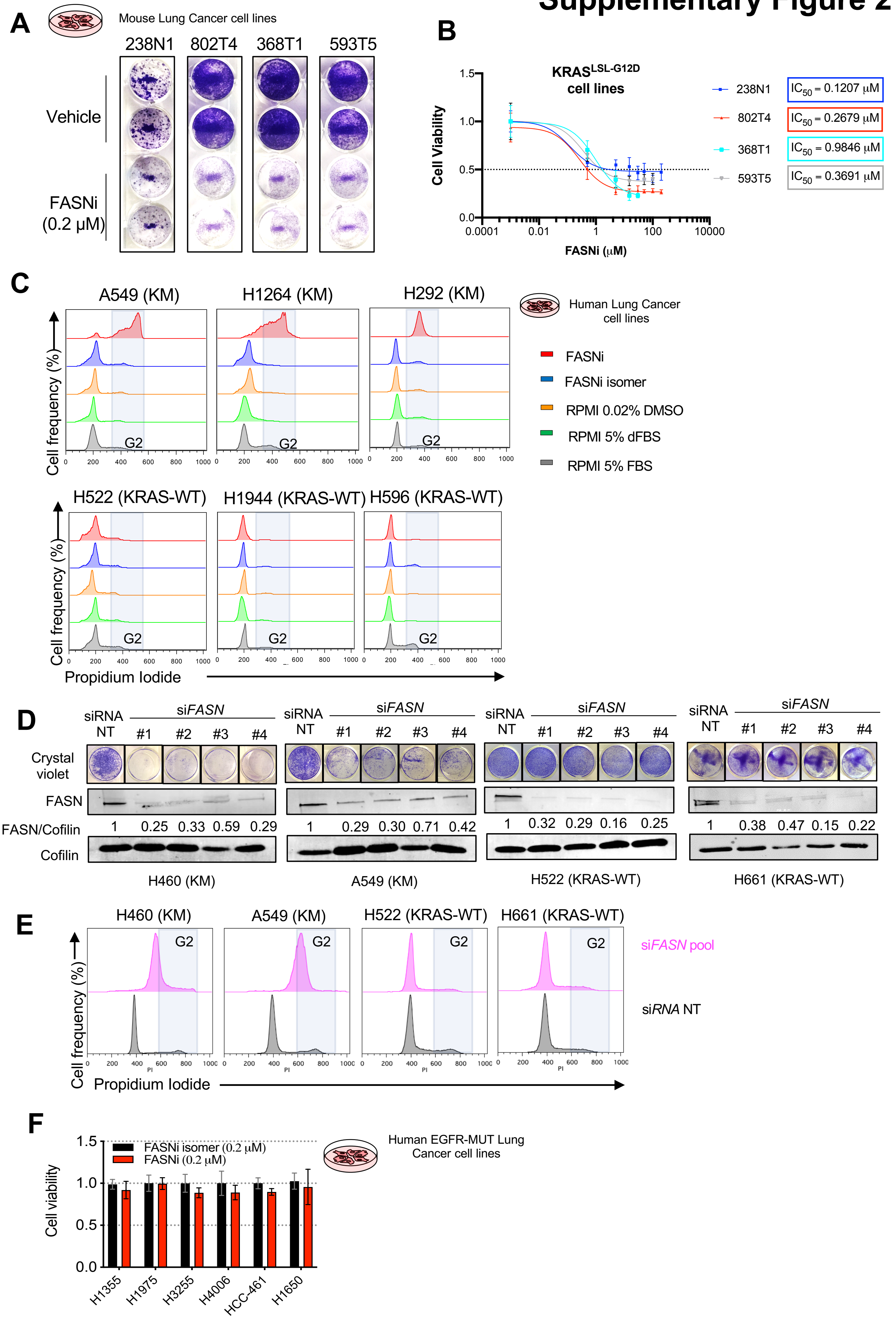

### Supplementary Figure 3

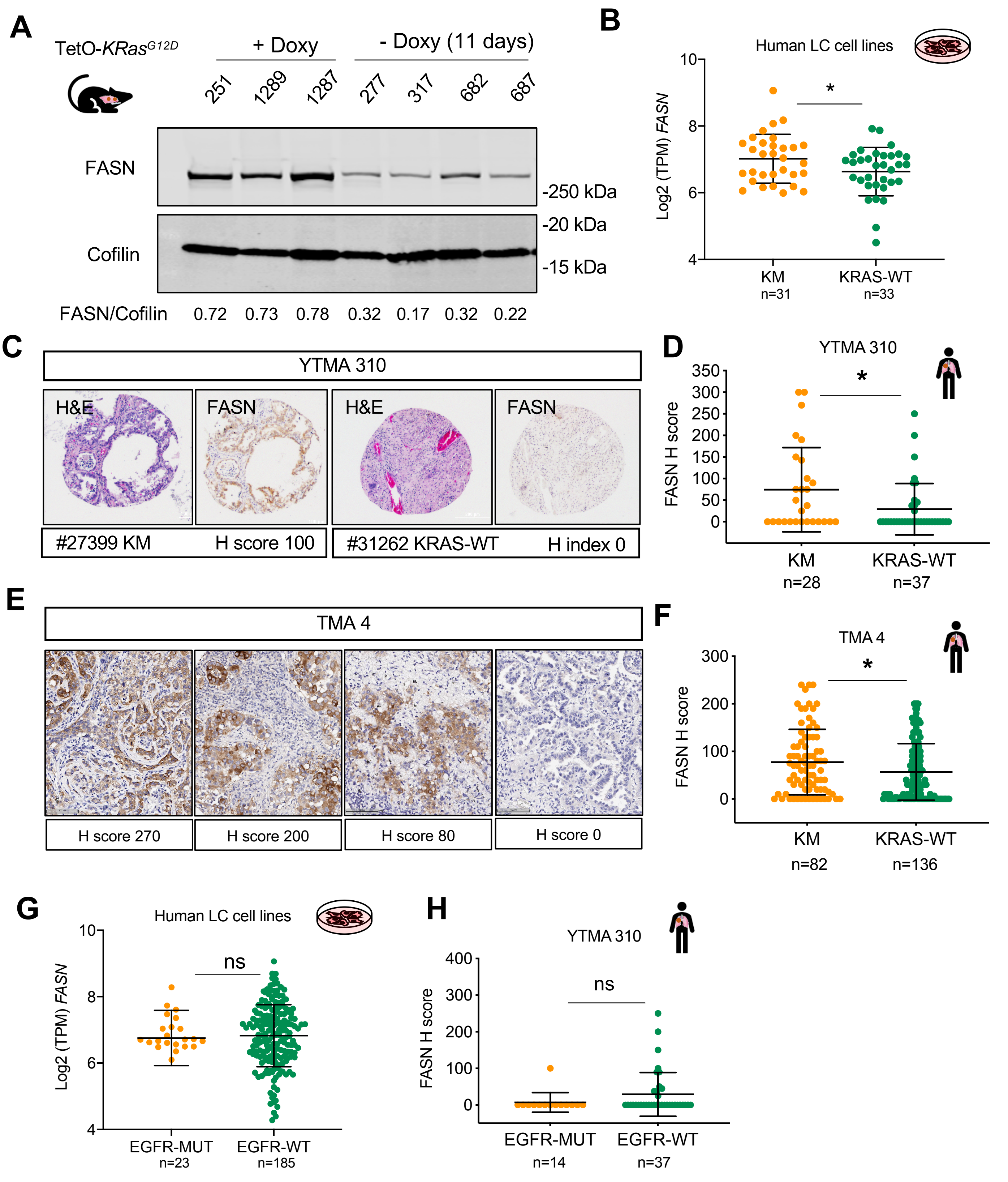

### Supplementary Figure 4

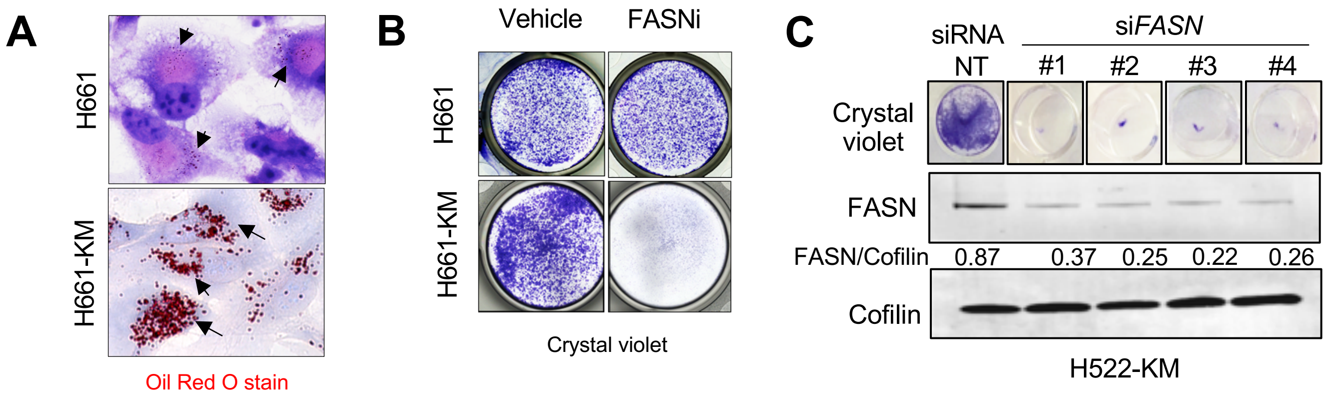

### Supplementary Figure 5

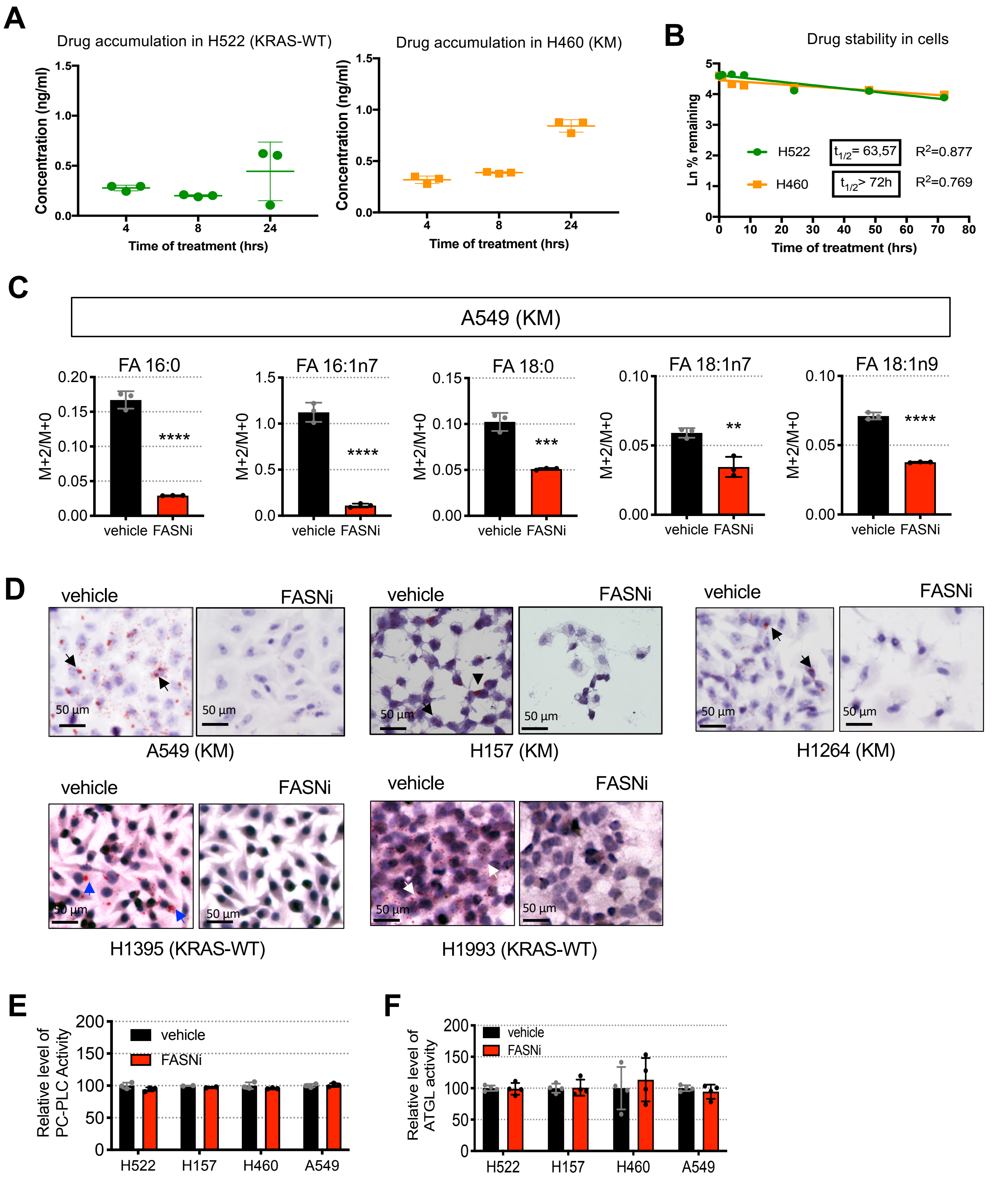

### Supplementary Figure 6

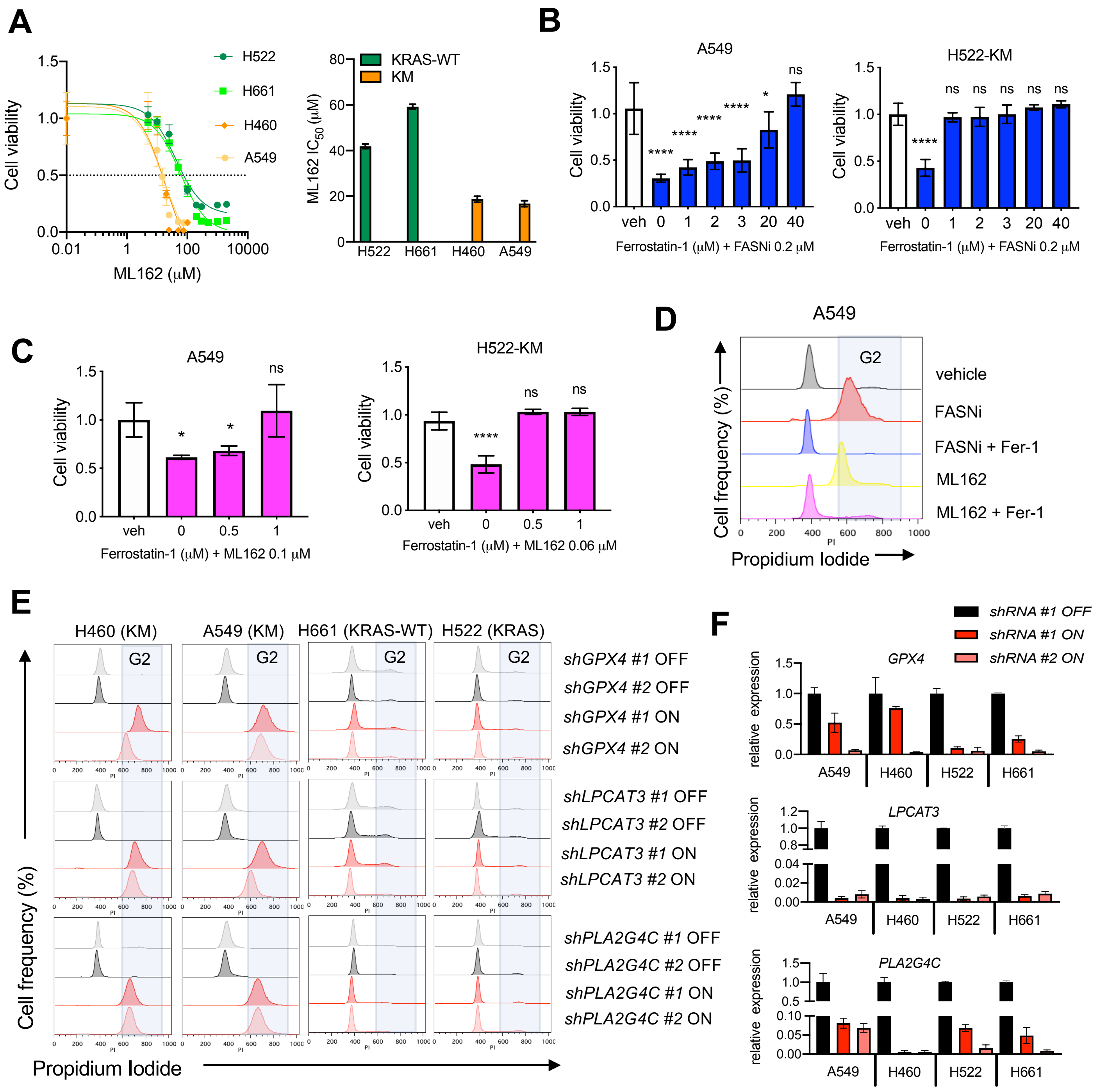

### Supplementary Figure 7

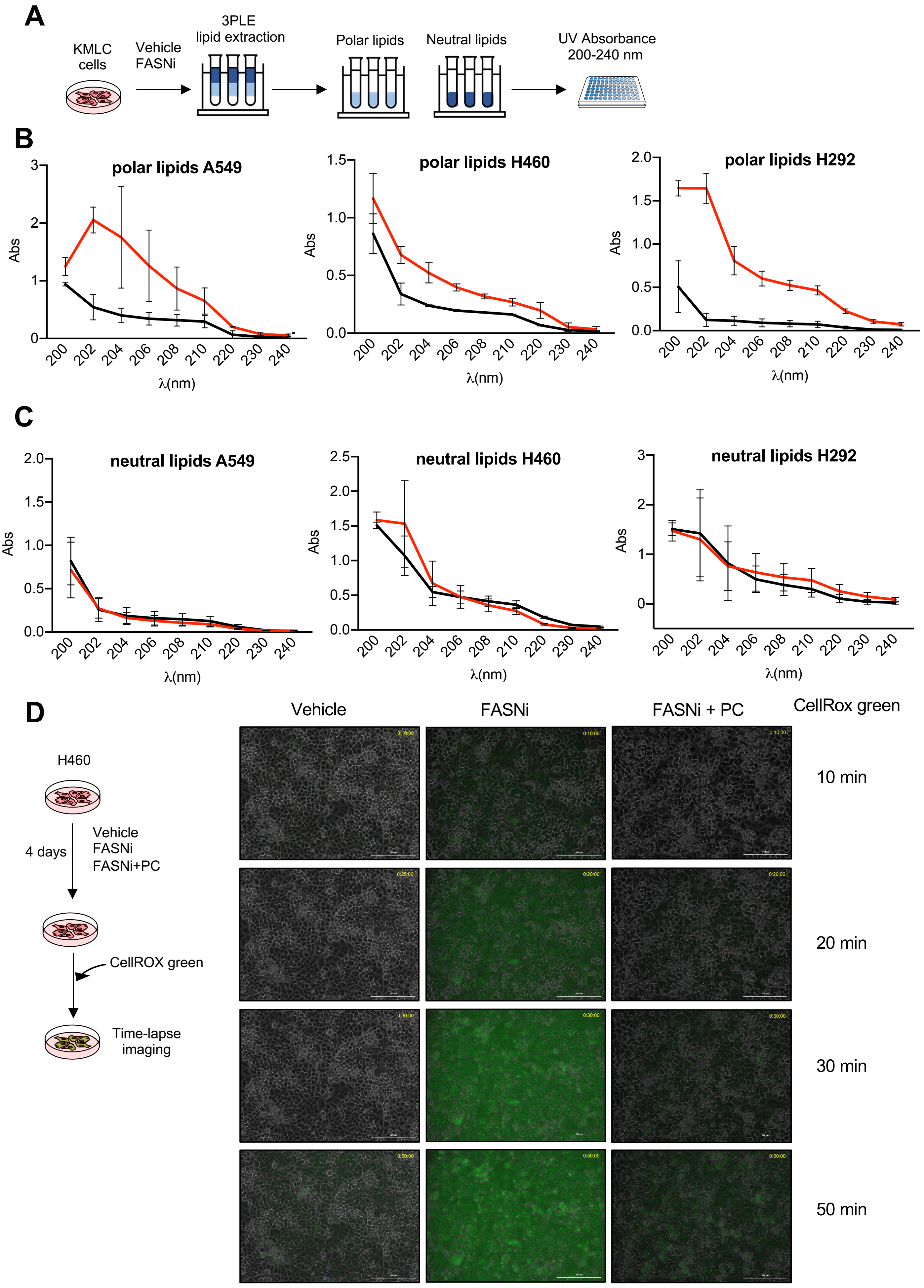

### Supplementary Figure 8

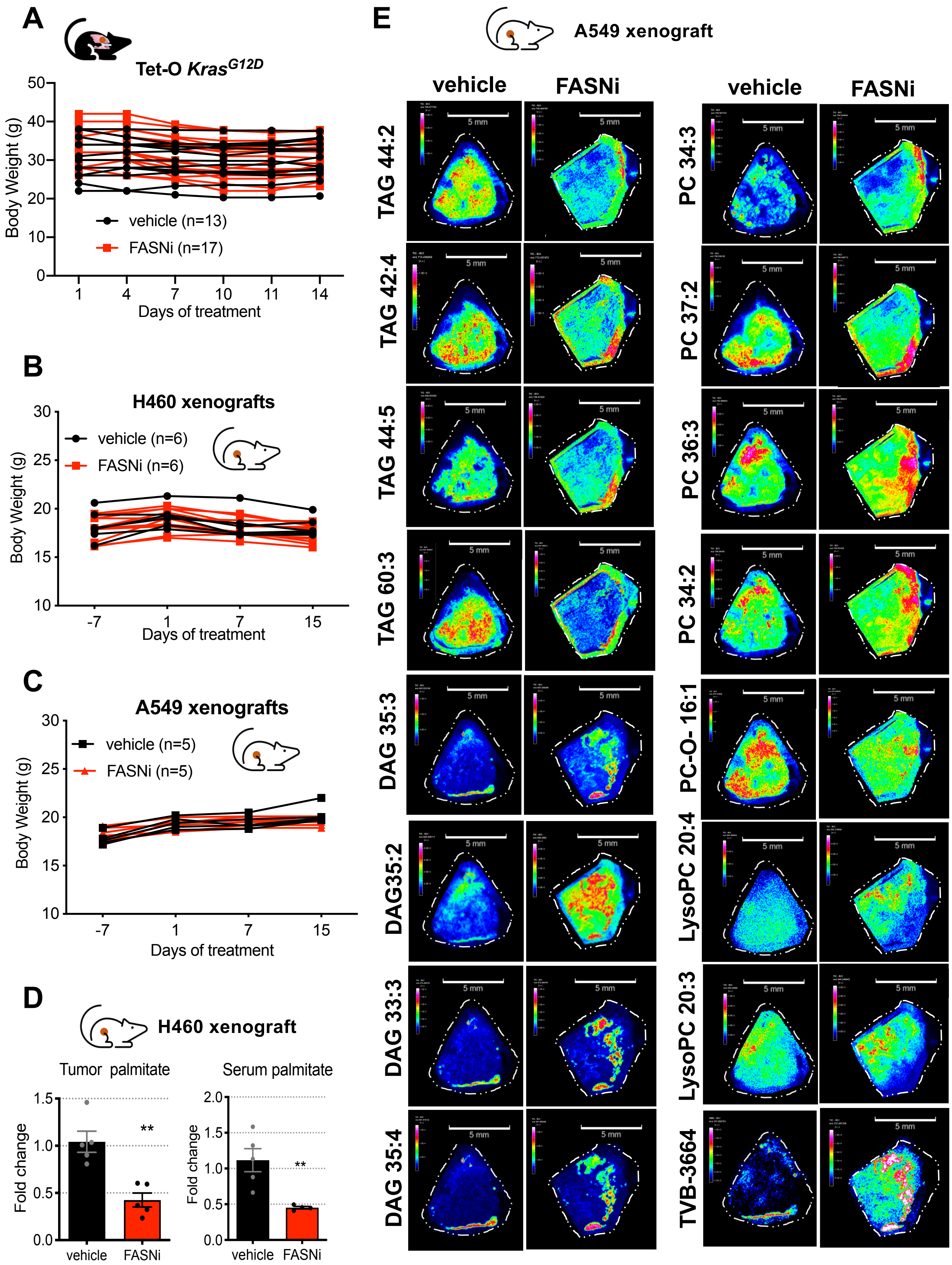
